# PhyClone: Accurate Bayesian reconstruction of cancer phylogenies from bulk sequencing

**DOI:** 10.1101/2024.08.14.607069

**Authors:** Emilia Hurtado, Alexandre Bouchard-Côté, Andrew Roth

## Abstract

**Motivation:** Cancer is driven by somatic mutations that result in the expansion of genomically distinct sub-populations of cells called clones. Identifying the clonal composition of tumours and understanding the evolutionary relationships between clones is a crucial task in cancer genomics. Bulk DNA sequencing is commonly used for studying the clonal composition of tumours, but it is challenging to infer the genetic relationship between different clones due to the mixture of different cell populations.

**Results:** In this work, we introduce a new probabilistic model called PhyClone that can infer clonal phylogenies from bulk sequencing data. We demonstrate the performance of PhyClone on simulated and real-world datasets and show that it outperforms previous methods in terms of accuracy and scalability.

**Availability and implementation:** Source code is available on Github at: https://github.com/Roth-Lab/PhyClone under the GPL v3.0 license.

## Introduction

Inferring the evolutionary history of cancer cell populations is a fundamental problem in cancer biology (Burrell et al., 2013; McGranahan and Swanton, 2017). High throughput genomics methods been transformative in this effort, providing insights into mechanisms of malignant ontogeny, therapeutic resistance and metastasis (Turajlic et al., 2019). While single-cell sequencing methods exist to precisely characterize tumour evolution (Zahn et al., 2017; Laks et al., 2019), their application has remained limited due to cost and technical challenges when applied to patient tumour samples. Thus, the field remains heavily reliant on bulk sequencing and computational deconvolution of tumours to infer clonal population structure (Dentro et al., 2021; Frankell et al., 2023).

Early tools for quantifying clonal populations in bulk sequencing focused on clustering mutations that occur in the same proportion of cancer cells (cellular prevalence) (Roth et al., 2014; Deveau et al., 2018; Gillis and Roth, 2020). These approaches assume that mutations with the same cellular prevalence across samples, share the same evolutionary history i.e. originate at the same point in evolutionary time and have the same pattern of mutation loss (Gerstung et al., 2020). While effective for identifying clonal populations, clustering approaches ignore the underlying phylogenetic relationship among clones. Subsequent approaches have attempted to reconstruct the clonal phylogeny from bulk sequencing (Deshwar et al., 2015; Malikic et al., 2015; Popic et al., 2015; Wintersinger et al., 2022; Grigoriadis et al., 2023b). Beyond providing an estimate of the phylogenetic tree, a key benefit of these approaches is that they allow for the genotypes and prevalence of clones to be resolved.

Methods for inferring clonal phylogenies from bulk sequencing generally assume that mutations with higher cellular prevalence occur earlier (closer to the root of the phylogeny) than mutations with lower cellular prevalence (McGranahan and Swanton, 2017). A corollary of this assumption is that if the cellular prevalence of two populations exceeds one, the lower cellular prevalence population must be a descendant of the higher cellular prevalence population (Dentro et al., 2017; Tarabichi et al., 2021). This additivity assumption is relatively weak, and cannot for example resolve whether two clonal populations are siblings or ancestor/descendants if the sum of cellular prevalence does not exceed one. This deficiency can partially be addressed by conducting multi-region sequencing (Tarabichi et al., 2021; Cortés-Ciriano et al., 2022), which has now become common practice in bulk cancer evolution studies (Frankell et al., 2023). The key insight is that if the ordering of cellular prevalence values between two populations switches in different samples, the clonal populations must be siblings (Tarabichi et al., 2021). In addition to only providing weak identifiability, the additivity assumption is violated if mutation loss occurs. In the context of cancer evolution, mutation loss can occur due to genomic instability leading to the deletion of large regions of the genome (McGranahan and Swanton, 2017; Tarabichi et al., 2021). Frequent mutation loss has been identified in genomically unstable cancers in several studies (McPherson et al., 2016; Jamal-Hanjani et al., 2017; Gerstung et al., 2020; Watkins et al., 2020).

In this work we develop a new method for inferring clonal phylogenies based on bulk sequencing. Given that bulk sequencing may only contain information to partially resolve the clonal phylogenetic structure, we adopt a Bayesian statistical approach which allows us to infer a posterior distribution over the set of potential phylogenies compatible with the data. To address the fact the number of clonal populations is unknown, we have developed a non-parametric Bayesian model of clusters related by a tree structure. A key benefit of our approach is the ability to marginalise the parameters associated with tree nodes (clonal prevalence), allowing us to perform posterior inference in the lower dimensional space of tree topologies. Inference in the collapsed space of discrete clustering trees still requires exploration of an exponential number of tree topologies with variable numbers of nodes. To address this issue, we developed an inference algorithm using particle Gibbs sampling (Andrieu et al., 2010) coupled with an auxiliary variable construction. In addition, our approach allows for partial information about clustering generated by fast non-phylogenetic methods to be incorporated. We demonstrate in the results, that these advances allow our approach to scale favourably in terms of running time and accuracy with increasing numbers of samples. In addition, we show how our proposed method can be adapted to allow for the potential of mutation loss for accurate inference in real datasets.

## Methods

The PhyClone model takes as input: i) allelic count data from somatic single nucleotide variants (SNVs); ii) copy number variant (CNV) information; iii) tumour content estimates from bulk sequence data of one or more related tumour samples (Figure 1). Using this information PhyClone infers the posterior distribution of phylogenetic trees relating the clones in the sample. As part of this inference the model infers the number of clones, sets of mutations which originate at each node in the tree, and the phylogenetic ordering of mutations. We do not explicitly infer the prevalence of clones in each sample, as this variable is marginalised during inference. If needed, it can be reinstantiated via maximum a posteriori (MAP) estimation or sampling after tree inference.

**Fig. 1.**
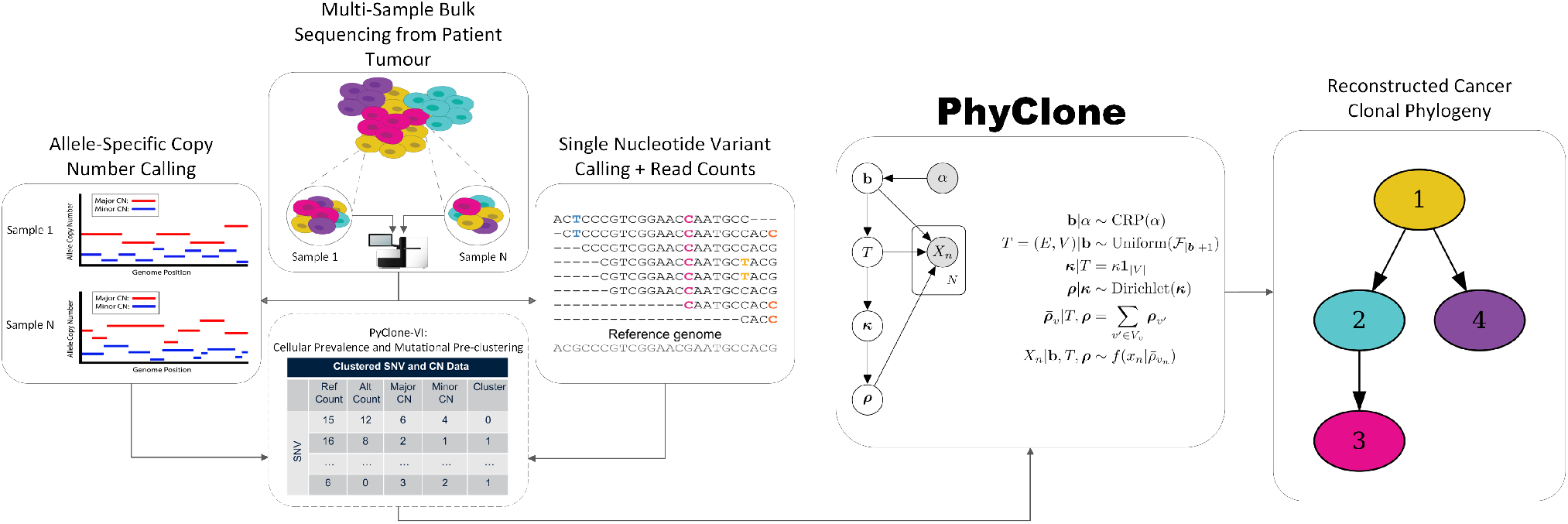
Overview of the PhyClone model. Outline of the high-level process of clonal phylogeny reconstruction from bulk sequencing used by PhyClone. As input PhyClone requires allele-specific read counts and copy-number data for single nucleotide variants (SNVs) identified in one or more samples.

We first describe the non-parametric Bayesian tree structured clustering prior distribution used by PhyClone. We then provide a full description of the basic PhyClone model, and several extensions. Finally, we describe the inference strategy. More detailed discussion of the model and inference strategy are provided in the Supplementary Methods.

### Prior distribution over clone phylogenies

PhyClone employs the forest structured Chinese restaurant process (FS-CRP) as a prior over tree topologies and clusters. In this section we briefly describe the generative process that defines the FS-CRP. In the first step of the FS-CRP we partition SNVs into clusters assumed to have the same evolutionary history. To do so SNVs are split into clusters via the Chinese restaurant process (CRP) (Aldous, 1985, pg. 92). In the second step of the FS-CRP, we sample a topology for the clonal phylogeny. To do so a multifurcating tree with as many nodes as clusters is selected uniformly at random and each cluster is associated to a node. The sampled tree may exhibit multiple roots, so we use the observation that any rooted forest (that is, any collection of rooted subtrees) over *K* nodes can be turned into a single rooted tree over *K* + 1 nodes by setting all root nodes to be children of a new empty root node. In the third step we assign clonal prevalence to each node. The clonal prevalence is defined to be the proportion of malignant cells which originate at a node, or equivalently the proportion of cells with the genotype defined by the union set of mutations assigned to nodes on the path to the root. To do so a vector of length equal to the number of nodes is sampled from a Dirichlet distribution. This enforces the constraint that the sum of all clonal prevalence adds to one. To obtain the cellular prevalence, that is the proportion of malignant cells that harbour an SNV, we move recursively up the tree from the leafs, summing the clonal prevalence of the node with the cellular prevalence of its children. This corresponds to assumption SNVs are propagated to all descendants from their clone of origin. The output of this process is a tree structured clustering of SNVs which we refer to as a clone phylogeny. In the clone phylogeny each node has a collection of originating SNVs, clonal prevalence, and cellular prevalence defined.

### PhyClone model

Figure 1 outlines the basic PhyClone model for a single sample with no outliers. The observed data vector *X* contains the allelic read counts and copy number information for all mutations *n* ∈ [*N*] = {1, …, *N*}. This model can be generalised to multi-region sequencing by sampling the clonal prevalence of each region from a Dirichlet distribution. To compute the cellular prevalence of a mutation, which will be used in the PyClone emission density for the allele counts of a mutation, let:

- *T*_*v*_ denote the subtree of *T* rooted at node *v*.
- *V*_*v*_ denote the set of nodes in *T*_*v*_.
- *C*_*v*_ denote the set of children nodes of *v*.
- *v*_*n*_ ∈ *V* denote the node associated with block *b* ∈ b such that *n* ∈ *b*, that is mutation *n* is in cluster *b*.
- 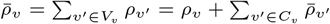

In this notation *ρ*_*v*_ and 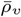 represents the clonal and cellular prevalence of node *v*.

With this notation the data likelihood is

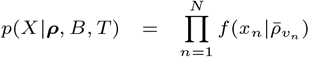

where 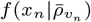 represents the copy number and tumour content corrected likelihood of allele counts based on the PyClone model (Supplementary Figure 1, Roth et al. (2014)).

#### Marginalising clonal prevalence

For inference, it is beneficial to work with the collapsed joint distribution, where we marginalise the node parameters ***ρ***. LetΔ_*k*_ denote the *k* simplex, 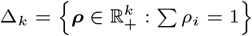.

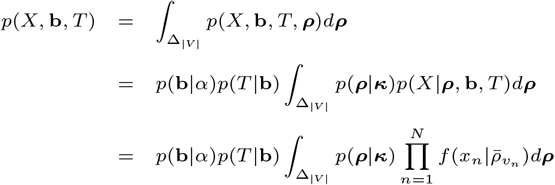

Computing 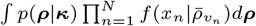 is non-trivial because of the dependence on the tree structure. We describe an efficient recursive algorithm in the Supplementary Material that computes the marginalised likelihood in *𝒪*(|*V* |^2^).

#### Pre-clustering

For computational efficiency, PhyClone is equipped to leverage clustering information from fast non-phylogenetic approaches such as PyClone-VI (Gillis and Roth, 2020).

Our approach enforces that any mutations clustered together by the pre-clustering step will be kept together. However, mutations assigned to different clusters by the pre-clustering algorithm can also be placed together by PhyClone. This allows PhyClone to refine the clustering based on the constraints imposed by the phylogeny. Though we did not make use of this in the experiments, it is theoretically beneficial for PhyClone analyses to run pre-clustering to overcluster the data i.e. generate more clusters than truth. PhyClone can then merge the resulting clusters to obtain the correct clustering.

#### Outliers

To accommodate SNVs which do not satisfy the additivity assumption due to mutation loss or sequencing error, we allow the possibility of outliers.

Each SNV is assigned a prior probability of being an outlier, *ν*_*n*_. We define a binary variable *o*_*n*_ which indicates whether mutation *n* is an outlier. If a mutation is not an outlier, it joins a node in the tree and contributes to the likelihood as above. If the mutation is an outlier, it has the standard PyClone likelihood where we assume it has a cellular prevalence drawn from a Uniform distribution. We marginalise the cellular prevalence in this case using numerical integration.

The joint likelihood with outliers becomes:

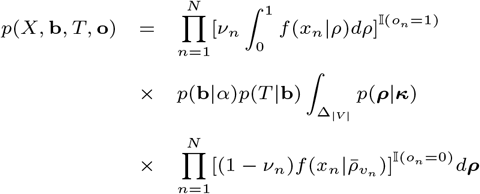

### Inference

To simultaneously infer the posterior distribution of tree topologies and cluster assignments (*T* and b), we use a sequential Monte Carlo (SMC) algorithm. The SMC algorithm works by evolving a set of particles which are iteratively extended and reweighed, such that the final iteration yields an approximation of the posterior.

We assume that a permutation, *σ*, of the mutation indices [*N*] has been given. At algorithmic time *t* of the SMC procedure we add data point *X*_*σ*(*t*)_ = *x*_*t*_.

A key parameter to define for SMC algorithms is the proposal function, *q*(*x*_*t*+1_|*x*_*t*_), which extends the state *x*_*t*_ to a new state *x*_*t*+1_. In the case of our algorithm, this corresponds to choosing which node in the tree to allocate mutation *σ*(*t* + 1) to. To propose a new state at time *t* + 1, we consider all possible states which can be reached by:

1. Adding a data point to an existing root node.
2. Creating a new node choosing a possibly empty subset of children from the existing root nodes as children.

We randomly choose whether to join an existing root node or start a new node. When adding a mutation to an existing root node, we compute the probabilities of the resulting trees with the mutation attached at each root node. When adding a mutation to a new node, we randomly sample the set of children. This approach provides a compromise between the quality of trees proposed and the computational complexity.

#### Overcoming the impact of data ordering

A limitation of the procedure so far described is that not all trees can be sampled due to the fixed order of the data points. Specifically, if *x* = *σ*_*i*_ and *y* = *σ*_*j*_ where *i* < *j* then any tree where *x* is an ancestor of *y* cannot be sampled given a fixed *σ* (illustrated in Supplementary Figures 2–4). For example, the first element of *σ* can never be a root node unless the sampled tree has only a single node. To address this issue we use a particle Gibbs (PG) sampler to embed our SMC procedure in a more general Markov chain Monte Carlo sampler (Andrieu et al., 2010).

At a high level, we treat *σ* as an auxiliary variable and alternate between sampling *σ* and (*T*, ***b***). Sampling of *σ* is constrained such that the SMC procedure could generate the current configuration of (*T*, ***b***).

To see that this algorithm yields a sampler that can sample all trees, note that the single node tree can be reached at any iteration of the algorithm. From this configuration, the sampling *σ* is uniform, thus allowing any ordering of mutations to be chosen for the subsequent update of (*T*, ***b***).

#### Additional MCMC moves

To improve the mixing of the sampler, it is beneficial to include additional moves in addition to the particle Gibbs updates. We use two moves, neither of which change the number of nodes in the tree.

The first move is a subtree prune-regraph. To perform this move, we select a node at random from the tree and remove the subtree rooted at this node. We then compute the probability of the tree which would result from attaching the subtree to a node in the tree. We do this for all nodes in the tree and Gibbs sample a new attachment node.

The second move we consider is to sample new node assignments for each data point in the tree without changing the topology. To do this, we first shuffle the data point ordering. Then, for each data point, if removing it from its current node does not empty the node, we compute the probability of each tree resulting from having added the data point to every node other than its currently assigned one. We use these probabilities to construct a probability distribution from which we then Gibbs sample the new tree from.

### Experiments

We tested PhyClone against three widely used state-of-the-art methods: CONIPHER (Grigoriadis et al., 2023b), Pairtree (Wintersinger et al., 2022), and PhyloWGS (Deshwar et al., 2015). Here we describe the datasets, program settings and performance metrics used for these benchmarking experiments.

#### Datasets

A combination of newly simulated synthetic datasets, previously published synthetic datasets, and real human cancer data were used to perform benchmarking of the methods.

##### Tree Structured Stick Breaking Data

Tree topologies and clusterings were simulated using the tree structured stick breaking (TSSB) prior. The TSSB prior is an alternative non-parametric Bayesian prior over clusters and tree topologies (Ghahramani et al., 2010). Allelic count data was simulated as diploid heterozygous mutations with a tumour content of 1.0. Two TSSB datasets were produced. Both datasets were simulated had a read depth of 1000, sample number levels of [2, 4, 8, 16], and tumour content of 1.0. The small dataset was simulated with 100 SNVs and the larger had 10,000 SNVs. Additionally, the small TSSB dataset also included runs with 128 samples.

##### Pairtree/Pairsim Data

We used previously published data from Wintersinger et al. (2022). The data was split into two groupings, low and high complexity, to reflect the difference in number of mutations, samples, and true tumour clusters/clones involved in each set of experiments. High complexity trials all varied according to: number of clusters, K={3, 10, 30, 100}; number of samples, S={10, 30, 100}; total depth, T={50, 200, 1000}; and number of mutations, M={30, 60, 100, 200, 300, 600, 1000, 2000, 3000, 10000} with four replicates each for a total of 432 trials. While the low complexity trials varied according to: number of clusters, K={3, 10}; number of samples, S={1, 3, 10}; total depth, T={50, 200, 1000}; and number of mutations, M={30, 60, 100, 200, 300, 1000} - with four replicates each for a total of 216 trials.

##### CONIPHER Data

We used previously published data from the Grigoriadis et al. (2023b,a). We used datasets 1 and 2, each with a total of 150 trials, which we refer to as noise and no-noise datasets respectively. For analysis, we split each dataset into three categories according to number of samples: 2–3, 4–7, and 8+.

##### FS-CRP loss data

Tree topologies and clusterings were simulated using the FS-CRP prior. Read count and copy-number data were simulated using the same method as the TSSB datasets. Mutational loss was simulated by first randomly selecting a set of mutations equal to the proportion to be lost, a node is then selected from which this set of mutations is to be ‘lost’ from; the loss-node is selected such that it is a descendant node from the node that the lost mutations originate from. All mutations that are members of the lost set are given positions on the same chromosome, to simulate the genomic locality of subclonal chromosomal region loss. All nodes downstream of (and including) the loss-node then have the cellular prevalence of the lost set of mutations subtracted from the node’s total, to reflect the impact of the having lost the mutations.

We simulated 100 trials for each level of mutation proportion to lose, with the mutation proportions being [0.0, 0.1, 0.2]. Each trial simulated 600 SNVs, with a depth of 1000, 8 samples, and a tumour content of 1.0.

##### HGSOC Data

Data used were from a previous single-cell study of High Grade Serous Ovarian Cancer (HGSOC) where ground truth trees were inferred from single-cell, whole genome, and targeted deep sequencing data (McPherson et al., 2016). Two sets of data were reconstructed for the HGSOC experiments; one dataset consisted of purely whole genome sequencing (WGS) bulk data, while the second dataset replaced the WGS allele counts with the targeted deep sequencing counts for the SNVs which were present in both sets. Pre-processing involved reconstruction of copy-number, read count, and cluster assignment data from the supplementary tables provided with the original study. Method inferred trees were constructed with the total set of processed SNVs per patient, and were then subset down to include only those SNVs that were used to define the ground truth trees from the original publication when computing metrics.

#### Program Parameters

Each method was allowed 48-hours wall-clock time to complete each individual trial in each dataset. Pre-clustering was performed for synthetic data using PyClone-VI initialised with 100 clusters, and 100 random restarts. For the HGSOC data, we used the clusters provided in the original publication. Pre-clustered results were supplied to CONIPHER, Pairtree and PhyClone for all experiments.

PhyClone (version 0.5.0) was run with the following parameters: 4 independent chains, 100 burn-in iterations, 5000 MCMC iterations, Beta-Binomial allele-count density, semi-adapted proposal kernel, final output tree selected by MAP joint-likelihood; and an outlier prior probability of 0.0. For experiments where we consider outliers in the PhyClone model, denoted PhyClone-O, we set the ‘--assign-loss-prob’ flag and accompanying outlier prior probabilities of 0.05 and 0.5 for the low and high loss prior probabilities respectively.

PhyloWGS, Pairtree and CONIPHER methods were run with default parameters, with the exception of CONIPHER in the HGSOC experiments, where the ‘min cluster size’ option was set to 3.

#### Metrics

Each experimental method was measured against ground truth in order to assess the accuracy of clonal phylogeny reconstruction and single nucleotide variant (SNV) assignments. The following metrics were employed:

- V-measure: A metric for measuring clustering accuracy (Rosenberg and Hirschberg, 2007)
- Ancestor-Descendant F-Score (AD F-Score): A reconstruction accuracy metric which measures the F1-score of a method’s phylogenetic reconstruction from the perspective of SNV ancestor-descendant relationships (Supplementary Figure 5). Positive cases result when an SNV is assigned to a node that can form an upward path to the node where the corresponding ancestor SNV is assigned.

Significance testing for differences in performance for each benchmarking experiment was performed via Friedman-Nemenyi testing. If the metric being tested demonstrated a global significance difference (*p*-value < 0.01) as per the Friedman test, a pairwise post-hoc Nemenyi test was then run to assess which methods performed significantly (*p*-value < 0.01) differently from each other. Subsequently, the mean difference was used to determine the direction of said significance (that is, which performed better out of the pair) (Demšar, 2006).

### Results

### Scalable and accurate Bayesian inference

We first analysed the small TSSB dataset with 100 SNVs. We note, this model is equivalent to the PhyloWGS model and should favour that method. Our goal was to compare the accuracy and efficiency of different modelling and inference approaches when data is generated from a well-defined statistical model. PhyClone and PhyloWGS are fully Bayesian models, whereas Pairtree uses a Gaussian approximation to perform MAP estimation, and CONIPHER optimises a heuristic objective. In addition, PhyClone, Pairtree and CONIPHER are all capable of utilizing pre-clustering of the data to scale inference.

The clustering and tree accuracy results for this experiment are summarised in Figure 2. PhyClone significantly outperformed all other methods in terms of V-measure and AD F-score. There was no significant difference in performance between Pairtree and PhyloWGS for either V-measure or AD F-score. CONIPHER was significantly outperformed by all methods in terms of both metrics. PhyloWGS was significantly slower than all other methods, requiring over an order of magnitude more time to finish than other approaches, with the gap widening with larger numbers of samples (Supplementary Figure 6).

**Fig. 2.**
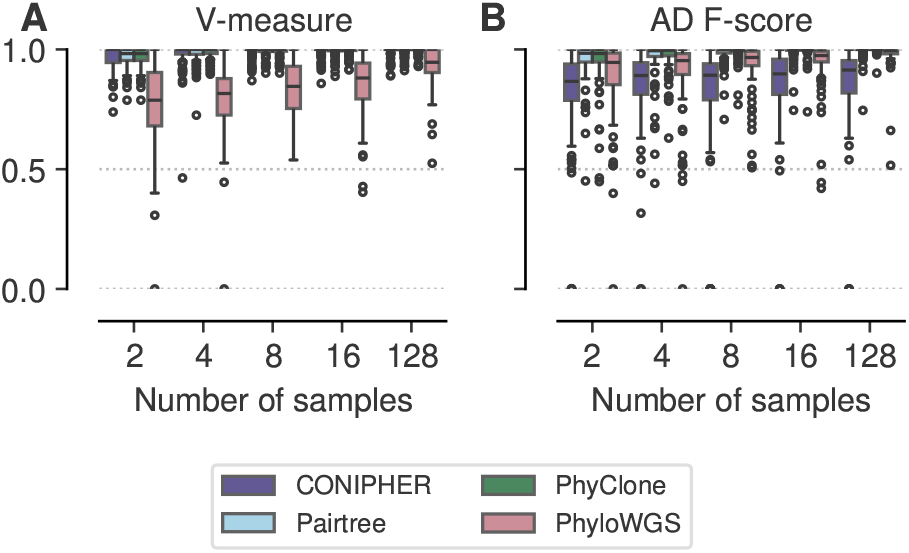
Performance on small TSSB dataset by number of samples. **A**. V-measure, clustering accuracy. **B**. Ancestor-Descendant F-Score.

This experiment supports the idea that fully Bayesian inference can be more accurate. Furthermore, pre-clustering of data can accelerate inference without compromising accuracy. Based on the long-running times observed for PhyloWGS, and the fact performance was not significantly better than other approaches under conditions which favour it, we did not include it in subsequent experiments.

### Robust performance across phylogenetic models

We next sought to compare the performance of the methods across a range of clonal phylogeny simulation strategies. We analysed the large TSSB, low and high complexity Pairtree (Pairtree-LC/HC) and CONIPHER no noise (CONIPHER-NN) datasets. Datasets spanned a range of copy number and tumour content values. All data in these experiments were simulated to satisfy the additivity constraint assumed by all models. Results for this experiment are summarised in Figure 3.

**Fig. 3.**
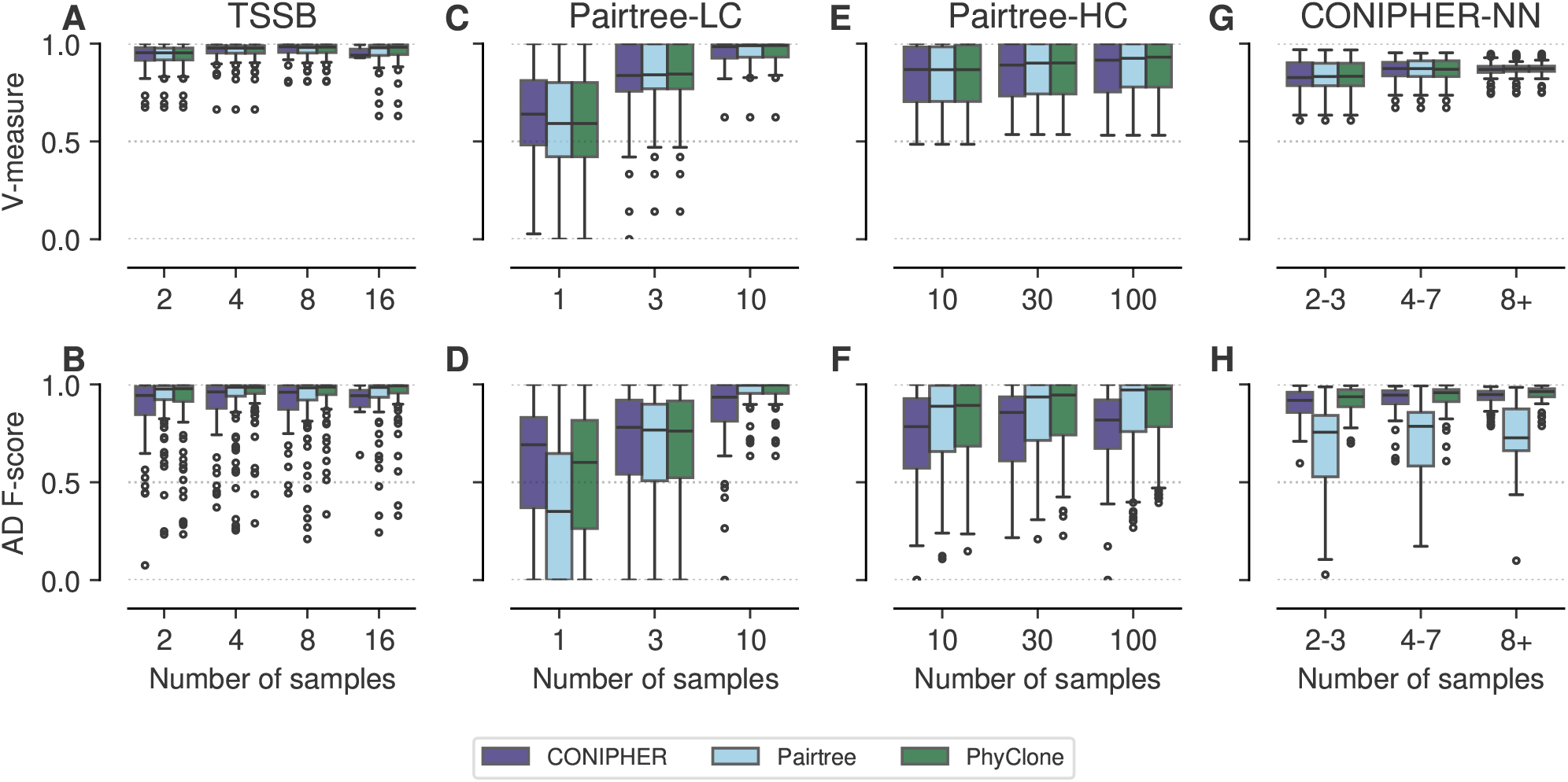
Performance on noise free synthetic data. V-measure and Ancestor-Descendant F-Score for **A-B** High SNV TSSB, **C-D** Pairsim low complexity, **E-F** Pairsim high complexity and **G-H** CONIPHER no noise datasets.

For both AD F-Score and V-Measure, PhyClone outperformed CONIPHER and Pairtree regardless of dataset. The performance advantages of PhyClone over Pairtree were significant for AD F-Score in the TSSB, Pairtree-HC, and CONIPHER-NN datasets; in terms of V-measure PhyClone significantly outperformed Pairtree in the TSSB dataset. The performance advantages of PhyClone over CONIPHER were significant for both metrics in all cases except for V-Measure on the CONIPHER-NN dataset, where there were no significant performance differences found between any method.

These experiments suggest that PhyClone is relatively robust to the phylogenetic data generation model. Furthermore, PhyClone tends to benefit more than other approaches from increasing numbers of samples. This is reflected in both increasing median performance, and reduction in interquartile range of results for both V-measure and AD F-score.

### Accurate inference in the presence of noise

We next considered the impact of “noise”, that is mutations which fail to satisfy the additivity constraint. We analysed a previously published data, which we refer to as the CONIPHER noise dataset, which includes clusters of mutations which do not satisfy the additivity constraint. The authors of this dataset suggest such mutations could come from sequencing error, or mutation loss. The results of this experiment are summarised in Figure 4.

**Fig. 4.**
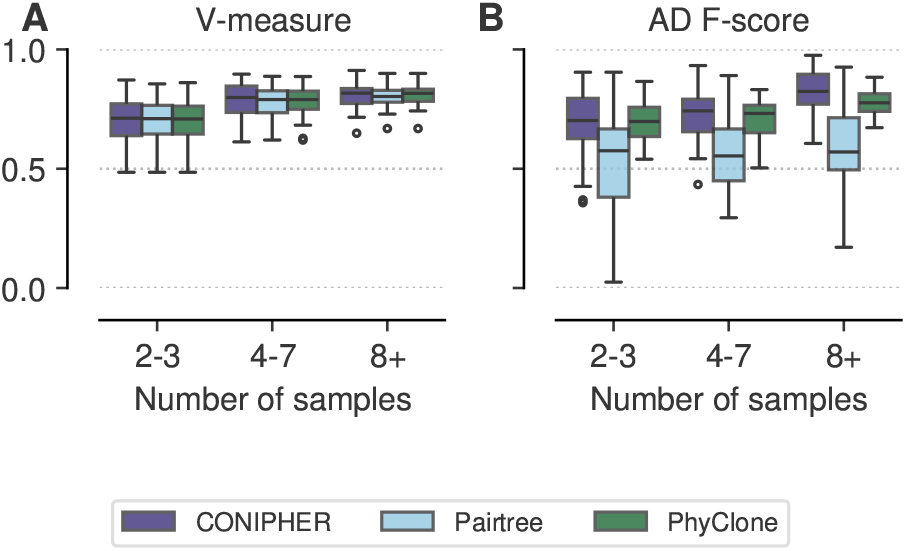
Performance on CONIPHER noise dataset. **A** V-measure and **B** Ancestor-Descendant F-Score for methods on the CONIPHER noise dataset.

No significant performance differences were found between any of the methods in terms of V-measure for this dataset. CONIPHER and PhyClone had no significant performance difference, and both significantly outperformed Pairtree in terms of AD F-score. Again, PhyClone benefited the most from increasing numbers of samples, with both better median performance and reduced variability.

### Outlier modelling mitigates additivity violation

The previous results suggest that PhyClone is generally robust to moderate violations of the additivity assumption, such as would be generated by sequencing error. However, larger violations of the additivity assumption, such as caused by the deletion of a large number of SNVs due to copy number variation could conceivably pose a challenge. To investigate this, we first simulated data from the PhyClone model allowing for mutation loss. We used this dataset to assess the performance of including an outlier state in the PhyClone model. As this dataset is inherently biased to favour PhyClone, we omit CONIPHER and Pairtree from this comparison. The results of this experiment suggest that the performance of the basic PhyClone model deteriorates as the proportion of mutations lost in the tree increases (Figure 5). However, the addition of the outlier state addresses this issue, making the method far more robust to loss.

**Fig. 5.**
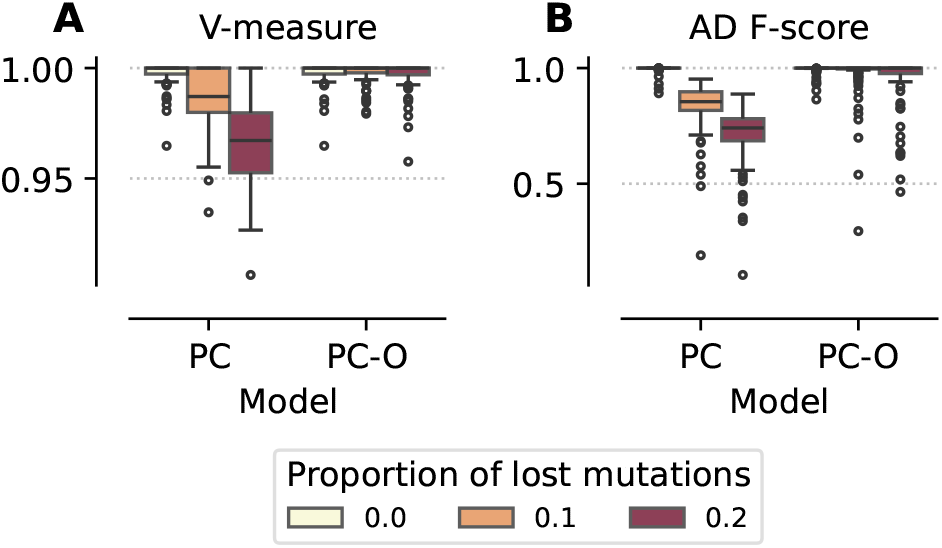
PhyClone performance with mutations loss. **A** V-measure and **B** Ancestor-Descendant F-Score for PhyClone with (PC-O) and without outlier modelling (PC) as a function of proportion of mutations lost.

### PhyClone allows for accurate inference of clone phylogenies with mutation loss

To assess how mutation loss impacts the performance of all methods, we analysed previously published data from three patients with high grade serous ovarian cancer (McPherson et al., 2016). Two patients (patients 2 and 3) exhibited complex forms of mutation loss, where clones experienced the loss of a subset of ancestral mutations. These results were validated with targeted single-cell sequencing, which we used as ground truth.

Pairtree struggled to resolve valid tree topologies (that is, a single-rooted directed acyclic graph) in most cases with this dataset, as such this section focuses on the results provided by PhyClone and CONIPHER.

PhyClone, PhyClone-O, and CONIPHER all appeared to capably reconstruct the phylogenetic history of the clones in patients 2 and 9, with varying levels of resolution (Supplementary Figures 7 and 8, Supplementary Table 1). For patient 3, in both forms of the data, WGS (Figure 6A) and WGS with targeted deep sequencing counts (Figure 6B), PhyClone-O was able to correctly identify and remove two SNV clusters that were found to be lost in ground truth clones below the root. Neither PhyClone without outlier modelling nor CONIPHER were able to remove the lost SNVs and thus were unable to completely reconstruct the ground truth phylogeny. The impact of the correct removal of the lost SNVs is further reflected in the performance metrics where PhyClone-O outperforms all methods for patient 3 (Supplementary Table 1).

**Fig. 6.**
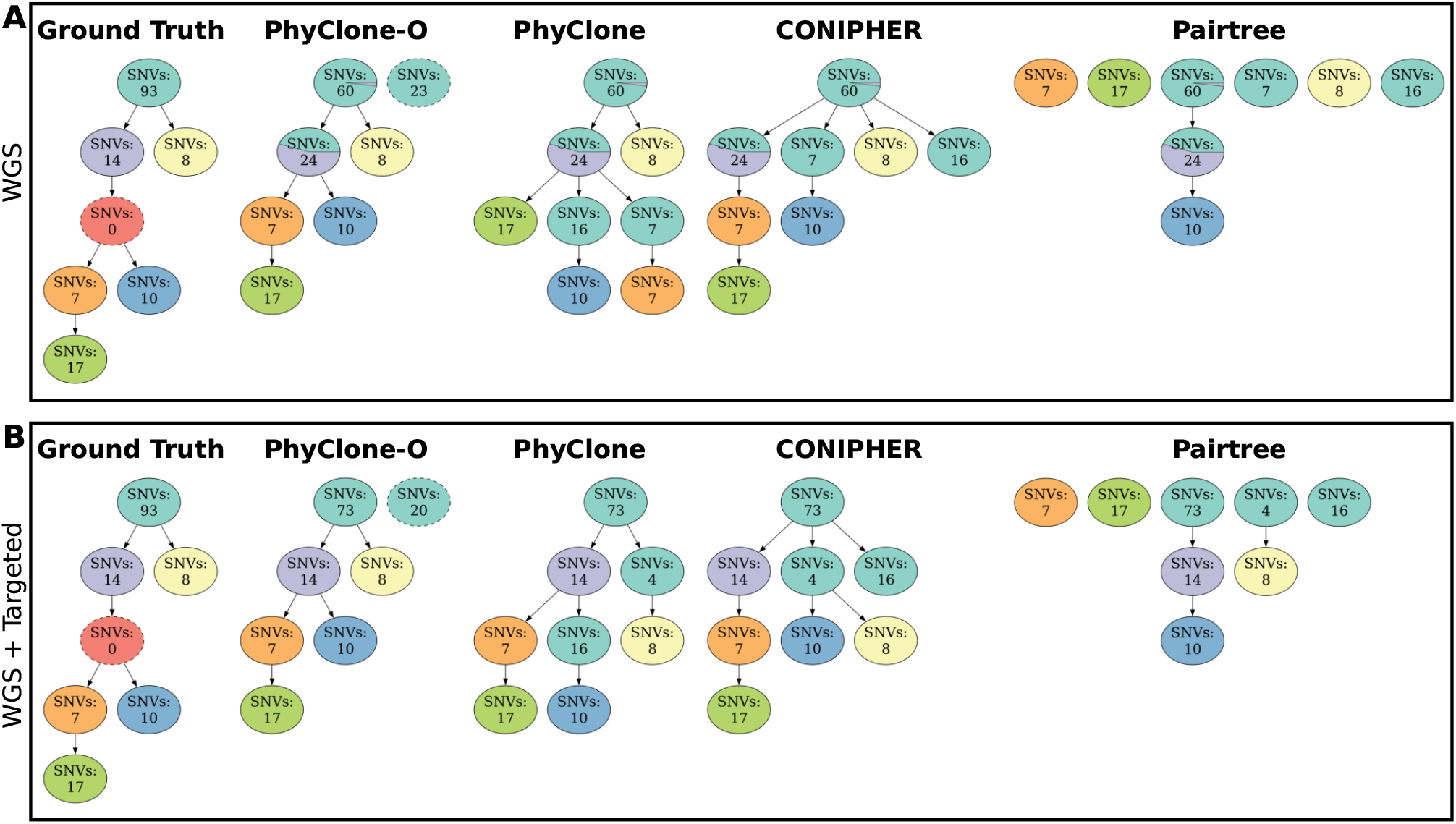
HGSOC patient 3 clonal phylogenetic trees. **A)** Predicted trees built using purely Whole Genome Sequencing (WGS) data. **B)** Predicted trees built using a combination of WGS and targeted deep sequencing data. (McPherson et al., 2016) (From left to ring): Ground truth phylogenetic tree inferred from single-cell and targeted deep sequencing data (McPherson et al., 2016), nodes with a dashed border denote clones that are defined only by the absence of SNVs from the parent. Predicted trees built using: PhyClone with outlier modelling; PhyClone without outlier modelling; CONIPHER; and Pairtree. Colours of nodes in method inferred trees correspond to the single nucleotide variant clonal (SNV) assignment from ground truth. Predicted tree nodes with a dashed border denote SNVs that the method has classified as outliers/lost.

## Discussion

In this work we have presented PhyClone, a new statistical method for reconstructing clonal phylogenies from multi-sample bulk sequencing data. PhyClone is based on a Bayesian non-parametric model, the FS-CRP, which we develop in this work. A key feature of using the FS-CRP in PhyClone, is the ability to marginalise the clonal prevalence given the constraints imposed by the tree. This allows us to perform collapsed inference in a parameter space that does not depend on the number of samples. As we demonstrate in the results, this improves the efficiency of sampling and accuracy of results as additional samples are used. In addition, we have developed a Particle Gibbs sampler for efficient inference. A key innovation of this inference scheme is an auxiliary variable construction over the sequence of points visited during each sequential Monte Carlo round.

The key limitation of PhyClone is the underlying PyClone model for correcting for mutational genotype. In particular, the model is not capable of modelling regions of sub-clonal copy number variation. However, as the number of samples increases we expect this deficiency to be less acute. For example, the HGSOC dataset analysed has numerous of sub-clonal copy number events (McPherson et al., 2016, 2017). In future, the PyClone model could also be improved to adjust fo sub-clonal copy number variation following approaches such as those in Frankell et al. (2023).

PhyClone presents a robust and scalable Bayesian method for clonal history reconstruction from multi-sample bulk-sequencing cancer data. Regardless of the source of synthetic data, PhyClone performed as well or frequently better than other state-of-the-art methods. Furthermore, PhyClone was able to reconstruct accurate phylogenies in real datasets in the presence of mutation loss. With its ability to leverage pre-clustering information, PhyClone easily scales to whole genome scale datasets. PhyClone also scales well in the number of samples analysed in terms of running time and accuracy, allowing for extensive multi-region sequencing datasets to be analysed.

## Conflict of interests

No competing interest is declared.

## Funding

A.R. is a Michael Smith Health Research BC scholar [grant 18245]. We acknowledge generous funding support provided to A.R. by the BC Cancer Foundation. In addition, A.R. receives operating funds from the Natural Sciences and Engineering Research Council of Canada (grant RGPIN-2022–04378).

## Data availability

Pairtree (Wintersinger et al., 2022), CONIPHER (Grigoriadis et al., 2023b), and HGSOC (McPherson et al., 2016) datasets are available from their respective publications. Pairtree datasets were retrieved from https://github.com/morrislab/pairtree-experiments/tree/master/inputs/sims.smallalpha.pairtree and CONIPHER datasets were retrieved from Grigoriadis et al. (2023a). TSSB synthetic datasets, FS-CRP loss synthetic datasets, and input files used for all experiments are available from Zenodo, DOI: 10.5281/zenodo.13240566.

## Notes

https://doi.org/10.5281/zenodo.13240566

### Competing Interest Statement

The authors have declared no competing interest.

