## Supplemental Methods and Figures for "PhyClone: Accurate Bayesian reconstruction of cancer phylogenies from bulk sequencing"

### 1 Supplementary Methods

#### 1.1 Allele count mode

PhyClone will use the same approach for correcting mutational genotype and tumour content as PyClone [3]. In this section we will review this approach mainly to establish notation. The data for the model is allele counts from $N$ mutations from $S$ samples. For simplicity, we will suppress the index $n$ for the mutation and $s$ for the sample in this section. We will assume that each mutation divides the set of cells that were sequenced into three sub-populations (Figure 1).

1. The normal cell population consisting of cells with healthy germline genomes.
2. The reference cell population which consists of cancer cells without the mutation in question.
3. The variant cell population which consists of cancer cells with the mutation in question.

Let $\mathcal{G} = (A, B, AA, AB, \cdots)$ be the set of all genotype where $A$ and $B$ represent reference and variant alleles respectively. For example $AB$ would represent a heterozygous variant with total copy number 2. We assume the genotype of all cells within each sub-population is constant. Let $\mathbf{G} = (G_N, G_R, G_V) \in \mathcal{G}^3$ be a vector where the entries are the genotype of the normal, reference and variant populations respectively. Let $t$ be the proportion of cancer cells in the sample. This is often referred to as the tumour content, tumour purity or cellularity of the sample. Let $\bar{\rho}$ be the proportion of cancer cells harbouring the mutation in the sample, that is the relative proportion of cancer cells in the variant population. This is often referred to as the cancer cell fraction (CCF) or cellular prevalence of the mutation. In sequel we will use tumour content and cellular prevalence.

Let $\epsilon$ be the assumed sequencing error rate. Let

1. $a(G) : \mathcal{G} \to \mathbb{N}$, $b(G) : \mathcal{G} \to \mathbb{N}$ be functions which map a genotype to the number of A and B alleles respectively.
2. $c(G) : \mathcal{G} \to \mathbb{N}$ be defined as $c(G) = a(G) + b(G)$ which is the total copy number of the loci.
3. $\mu(G) : \mathcal{G} \to \mathbb{N}$ be defined as:

$$\mu(G) = \min\left\{\max\left\{\frac{b(G)}{c(G)}, \epsilon\right\}, 1-\epsilon\right\}$$

   Which can be interpreted as the probability of sampling a read with the mutation from a population with genotype $G$.

Let $\xi(\mathbf{G}, \bar{\rho}, t)$ be the probability of sampling a read with the variant allele. We assume that we have an infinite initial population of cells which are sampled when sequencing. With this assumption the probability of sampling a read with a variant allele is roughly proportional to the number of copies of the variant allele in the input pool of DNA. More formally, accounting for sequencing error, the probability of sampling a variant allele is given by the following equation.

$$\begin{aligned}\xi(\mathbf{G}, \bar{\rho}, t) &= \frac{1}{Z}(1-t)c(G_N)\mu(G_N) \\ &+ \frac{1}{Z}t(1-\bar{\rho})c(G_R)\mu(G_R) \\ &+ \frac{1}{Z}t\bar{\rho}c(G_V)\mu(G_V) \\ Z &= (1-t)c(G_N) + t(1-\bar{\rho})c(G_R) + t\bar{\rho}c(G_V)\end{aligned}$$

Now we observe $d$ total reads covering the mutation in the sample, of which $x$ contain the mutant allele. Thus

$$p(x|d, \boldsymbol{G}, \bar{\rho}, t) = \text{Binomial}(x|d, \xi(\mathbf{G}, \bar{\rho}, t))$$

If the data has more variance than can be explained by a Binomial model we can instead use

$$p(x|d, \boldsymbol{G}, \bar{\rho}, t, \gamma) = \text{BetaBinomial}(x|d, \xi(\mathbf{G}, \bar{\rho}, t), \gamma)$$

where the Beta-binomial distribution is parameterised by the mean $\xi(\mathbf{G}, \bar{\rho}, t)$ and precision (inverse of variance) $\gamma$.

So far we have assumed the genotypes of the sub-populations were known. In general this is not true for the reference and the variant populations. Instead it is typical to observe allele specific copy number estimates for the region overlapping a mutation. Using this information we can elicit a prior over a set of plausible genotypes. We explain how to do this in the next section. For now assume we have a vector $\boldsymbol{\pi}$ of prior probabilities where $\pi_i$ is the prior probability of the $i^{th}$ plausible joint genotype, $\mathbf{G}_i$, of the populations. We can write to the probability of the observed data marginalising over all plausible genotypes as follows.

$$p(x|d, \boldsymbol{\pi}, \bar{\rho}, t) = \sum_i \pi_i \text{Binomial}(x|d, \xi(\mathbf{G}_i, \bar{\rho}, t))$$

or

$$p(x|d, \boldsymbol{\pi}, \bar{\rho}, t, \gamma) = \sum_i \pi_i \text{BetaBinomial}(x|d, \xi(\mathbf{G}_i, \bar{\rho}, t), \gamma)$$

We will call this the PyClone$(x|d, \boldsymbol{\pi}, \bar{\rho}, t)$ distribution in sequel where with some abuse of notation we ignore whether a Binomial or Beta-Binomial distribution is used.

We note that if we define the clonal prevalence of a mutation to be the proportion of cancer cells from a clonal population. Then the cellular prevalence is the sum of clonal prevalence for all clonal populations which have the mutation, assuming mutations are not lost. In what follows the clonal prevalence will be the primary quantity of interest, though the cellular prevalence is required for the allele count model.

#### 1.2 Eliciting mutational genotype priors

Let $c_{major}$ and $c_{minor}$ denote the major and minor allele copy number for the region overlapping the mutation in the sample. We will use the "major copy number" method for setting potential genotype priors. This method considers two cases. In the first case, the mutation occurs before the copy number event. In this case the reference population genotype matches the normal population. We consider all possible mutational genotypes for the variant population with up to $c_{major}$ chromosomes containing the variant. In the second case, the mutation occurs after the copy number event. In this case the reference population has $c_{major} + c_{minor}$ reference alleles. The variant population has 1 variant allele and $c_{major} + c_{minor} - 1$ reference allele. We set the prior weights to be equal for all possible mutational genotypes. For example suppose we have that $c_{major} = 2$ and $c_{minor} = 1$ and the normal copy number is 2. We have the following possible genotypes

- $\mathbf{G}_1 = (AA, AA, AAB)$
- $\mathbf{G}_2 = (AA, AA, ABB)$
- $\mathbf{G}_3 = (AA, AAA, AAB)$

each with prior probability $\frac{1}{3}$. Note that if allele specific copy number is not available then $c_{major}$ can be set to the total copy number and $c_{minor}$ to zero.

#### 1.3 Generative model

In what follows we let $F = (E, V)$ denote a rooted forest, that is a graph with a directed edge set $E$ and vertices set $V$. A root node is defined to be a vertex in $V$ with incoming degree zero. A rooted tree, denoted by $T = (E, V)$, is a rooted forest with one root node. We define $\mathcal{F}$ to be the space of all rooted forests, and $\mathcal{F}_k = \{F = (E, V) \in \mathcal{F} : |V| = k\}$ to be the set of all forests with $k$ nodes. We let $\mathbf{b} = \{b \subset \{1, \ldots, N\} : \bigsqcup b = [N]\}$ denote a partition of the data points.

The PhyClone generative model without outliers is as follows.

$$\begin{aligned}
\mathbf{b}|\alpha &\sim \text{CRP}(\alpha) \\
F = (E', V')|\mathbf{b} &\sim \text{Uniform}(\mathcal{F}_{|\boldsymbol{b}|}) \\
V|V' &= V \cup \{r\} \\
E|E' &= E' \cup \{(r, u) : \text{indegree}(u) = 0\} \\
T &= (E, V) \\
\boldsymbol{\kappa}|T &= \kappa \mathbf{1}_{|V|} \\
\boldsymbol{\rho}|\boldsymbol{\kappa} &\sim \text{Dirichlet}(\boldsymbol{\kappa})
\end{aligned}$$

where the notation $\mathbf{1}_k$ indicates the vector of ones with dimension $k$.

Thus, we sample a partition $\mathbf{b}$ from a CRP. Given $\mathbf{b}$ we sample a forest uniformly at random from the set of forests of size $|\mathbf{b}|$. We then deterministically connect all root nodes in the forest to a dummy node $r$ to create a tree. Next we sample a vector, $\boldsymbol{\rho}$, of clonal prevalences from a Dirichlet distribution of dimension $|\mathbf{b}| = |V|$. Each node is associated with a clonal prevalence in this vector. This model can be trivially generalized to multi-region sequencing by sampling the clonal prevalence of each region from a Dirichlet distribution. For simplicity we will assume a single sample in this section to keep the notation uncluttered. We also note that $\boldsymbol{\rho}$ depends on $T$ since the dimensionality of $\boldsymbol{\kappa}$ is determined by $|V|$. To simplify notation we do not write this dependence explicitly in some equations, but it should always be understood.

Next we define how to compute the cellular prevalence of a mutation, which will be used in the PyClone emission density for the allele counts of a mutation. Let

- $T_v$ denote the sub-tree of $T$ rooted at node $v$.
- $V_v$ denote the set of nodes in $T_v$.
- $C_v$ denote the set of children nodes of $v$.
- $v_n \in V$ denote the node associated with block $b \in \mathbf{b}$ such that $n \in b$, that is mutation $n$ is in cluster $b$.
- $\bar{\rho}_v = \sum_{v' \in V_v} \rho_{v'} = \rho_v + \sum_{v' \in C_v} \bar{\rho}_{v'}$

In this notation $\bar{\rho}_{v_n}$ represents the cellular prevalence of mutation $n$. With this notation the data likelihood is

$$p(X|\boldsymbol{\rho}, B, T) = \prod_{n=1}^{N} f(x_n|\bar{\rho}_{v_n})$$

where $f(x|\bar{\rho})$ is the PyClone emission density defined in supplementary section 1.1.

Thus, the joint density is given by

$$\begin{aligned} p(X, \mathbf{b}, T, \boldsymbol{\rho}) &= p(\mathbf{b}|\alpha)p(T|\mathbf{b})p(\boldsymbol{\rho}|\boldsymbol{\kappa})p(X|\boldsymbol{\rho}, \mathbf{b}, T) \\ p(\mathbf{b}|\alpha) &\propto \alpha^{|\mathbf{b}|} \prod_{b \in \mathbf{b}} (|b| - 1)! \\ p(F|\mathbf{b}) &= \frac{1}{(|\mathbf{b}| + 1)^{|\mathbf{b}|-1}} \end{aligned}$$

The normalisation of $p(F|\mathbf{b})$ can be derived from Cayley's formula for the number of rooted trees. We use the observation that any rooted forest over $K$ nodes can be turned into a rooted tree over $K + 1$ nodes by setting all root nodes to be children of a new dummy root node.

##### 1.3.1 Penalizing multi-rooted final tree topologies

One adjustment to the above is made when computing the prior density of the tree that follows after the addition of all datapoints has been completed. This adjustment was found to be necessary to address a bias towards trees with multiple children beneath the dummy node, referred to as sub-roots, which once removed would represent a multi-rooted tree. The adjustment itself is to penalize the likelihood of trees with more than one sub-root. Let the root adjustment term be $p(r|\mathbf{b})$, where the constant $C$ is chosen such that the single rooted tree is 1000 times more likely than the two root tree etc., and where $r$ is the number of sub-roots:

$$\begin{aligned} C &= 1000 \\ Z &= \sum_{i=1}^{|\boldsymbol{b}|} 1^{|\boldsymbol{b}|} \frac{1}{C^{i-1}} \\ p(r|\mathbf{b}) &= \frac{1}{Z * C^{r-1}} \end{aligned}$$

Let $w$ represent the number of ways that the current topology with its number of nodes, sub-roots, and sub-root topologies can exist; which is computed by applying Cayley's formula for the number of rooted trees on each of these sub-root trees and computing the product of the results. Let the set of sub-root trees be $\boldsymbol{S}$. The normalisation of $p(F|\mathbf{b})$ with the root adjustment term then becomes:

$$\begin{aligned} p(w|\mathbf{b}) &= \prod_{s_r \in \boldsymbol{S}} (|s_r|+1)^{|s_r|} \\ p(F|\mathbf{b}) &= \frac{1}{w} * p(r|\mathbf{b}) \end{aligned}$$

#### 1.4 Collapsed distribution

For inference it is beneficial to work with the collapsed joint distribution, where we marginalise the node parameters $\boldsymbol{\rho}$. Let $\Delta_k$ denote the $k$ simplex, $\Delta_k = \left\{\boldsymbol{\rho} \in \mathbb{R}_+^k : \sum \rho_i = 1\right\}$.

$$\begin{aligned} p(X, \mathbf{b}, T) &= \int_{\Delta_{|V|}} p(X, \mathbf{b}, T, \boldsymbol{\rho}) d\boldsymbol{\rho} \\ &= p(\mathbf{b}|\alpha) p(T|\mathbf{b}) \int_{\Delta_{|V|}} p(\boldsymbol{\rho}|\boldsymbol{\kappa}) p(X|\boldsymbol{\rho}, \mathbf{b}, T) d\boldsymbol{\rho} \\ &= p(\mathbf{b}|\alpha) p(T|\mathbf{b}) \int_{\Delta_{|V|}} p(\boldsymbol{\rho}|\boldsymbol{\kappa}) \prod_{n=1}^{N} f(x_n|\bar{\rho}_{v_n}) d\boldsymbol{\rho} \end{aligned}$$

Computing $\int p(\boldsymbol{\rho}|\boldsymbol{\kappa}) \prod_{n=1}^{N} f(x_n|\bar{\rho}_{v_n}) d\boldsymbol{\rho}$ is non-trivial because of the dependence on the tree structure.

We note that $\boldsymbol{\rho}$ has a Dirichlet distribution thus $p(\boldsymbol{\rho}|\boldsymbol{\kappa}) = c(\boldsymbol{\kappa}) \prod_{v \in V} \rho_v^{\kappa_v - 1}$. Thus, we can reorganise the computation. In the following we write $n \in v$ to denote that mutation $n$ is in the block $b \in \mathbf{b}$ associated with node $v$.

$$\begin{aligned} \int_{\Delta_{|V|}} p(\boldsymbol{\rho}|\boldsymbol{\kappa}) \prod_{n=1}^{N} f(x_n|\bar{\rho}_{v_n}) d\boldsymbol{\rho} &= \int_{\Delta_{|V|}} p(\boldsymbol{\rho}|T) \times \prod_{v \in V} \prod_{n \in v} f(x_n|\bar{\rho}_v) d\boldsymbol{\rho} \\ &\propto \int_{\Delta_{|V|}} \left[\prod_{v \in V} \rho_v^{\kappa_v - 1} \prod_{n \in v} f(x_n|\bar{\rho}_v)\right] d\boldsymbol{\rho} \\ &= \int_{\mathcal{S}_T} \left[\prod_{v \in V} \left(\bar{\rho}_v - \sum_{v' \in C_v} \bar{\rho}_{v'}\right)^{\kappa_v - 1} \prod_{n \in v} f(x_n|\bar{\rho}_v)\right] d\bar{\boldsymbol{\rho}} \end{aligned}$$

We perform a change of variable into the $\bar{\rho}_v$ parameterization and consequently also change the domain of integration into $\mathcal{S}_T = \left\{\bar{\boldsymbol{\rho}} \in \mathbb{R}_+^{|V|} : \sum_{v' \in C_v} \bar{\rho}_{v'} \leq \bar{\rho}_v \leq 1\right\}$.

To make further progress we use a discrete approximation for $\bar{\boldsymbol{\rho}} = (\bar{\rho}_1, \ldots, \bar{\rho}_{|V|})$. Specifically we assume that for a grid of size $L+1$, $\bar{\rho}_v \in \Phi = [0, \frac{1}{L}, \ldots, \frac{L-1}{L}, 1]$. We write $\mathcal{S}_{T,L}$ for the corresponding discretization of $\mathcal{S}_T$.

#### 1.5 Algorithm

Let $R_v(\bar{\rho})$ denote the likelihood of a sub-tree rooted at $v \in V$ when the mutations located at the root of this sub-tree have cellular prevalence $\bar{\rho} \in \Phi$. Let $\ell_v(\bar{\rho}_v) = \prod_{n \in v} f(x_n|\bar{\rho}_v)$. If we denote the sub-tree of $T$ rooted at $v$ by $T_v = (V_v, E_v)$, this has the form:

$$\begin{aligned} R_v(\bar{\rho}) &= \sum_{\bar{\boldsymbol{\rho}} \in \mathcal{S}_{T,L} : \bar{\rho}_v = \bar{\rho}} \prod_{v' \in V_v} {\rho_{v'}}^{\kappa - 1} \ell_{v'}(\bar{\rho}_{v'}) \\ &= \sum_{\bar{\boldsymbol{\rho}} \in \mathcal{S}_{T,L} : \bar{\rho}_v = \bar{\rho}} \prod_{v' \in V_v} \left(\bar{\rho}_{v'} - \sum_{v'' \in C_v} \bar{\rho}_{v''}\right)^{\kappa - 1} \ell_{v'}(\bar{\rho}_{v'}) \end{aligned}$$

We need to show that $R_v(\cdot)$ can be efficiently computed from the recursions $\{R_{v'}(\cdot) : v' \in C_v\}$. This is done in three steps. In this section, we show the steps taken by the algorithm, and we analyse and justify them in supplementary section 1.5.1. The pseudo-code for the marginalisation algorithm consists, for all $v \in V$ in post-order with respect to $T$:

1. A first sub-recursion, which iteratively incorporates the sub-recursions obtained for each of the $|C_v|$ children of node $v$:

$$\begin{aligned} D_v^1(\bar{\rho}) &= R_{v_1}(\bar{\rho}), \\ D_v^k(\bar{\rho}) &= \sum_{\bar{\rho}' \in \Phi : \bar{\rho}' \leq \bar{\rho}} R_{v_n}(\bar{\rho}) D_v^{k-1}(\bar{\rho} - \bar{\rho}'),\ k \in (2, \ldots, |C_v|), \end{aligned}$$

2. A second sub-recursion to marginalise the sum of the children's total prevalence, $\sum_k \bar{\rho}_{v_k} = \bar{\rho}'$, while taking into consideration the prior contribution based on the residual prevalence assigned to the current node $v$:

$$S_v(\bar{\rho}) = \sum_{\bar{\rho}' \in \Phi : \bar{\rho}' \leq \bar{\rho}} (\bar{\rho} - \bar{\rho}')^{\kappa - 1} D_v^{|C_v|}(\bar{\rho}'),$$

which, when $\kappa = 1$, reduces to:

$$\begin{aligned} S_v(0) &= 0, \\ S_v(\bar{\rho}) &= D_v^{|C_v|}(\bar{\rho}) + S_v\left(\bar{\rho} - \frac{1}{L}\right),\ \bar{\rho} \in \left(\frac{1}{L}, \ldots, \frac{L}{L}\right). \end{aligned}$$

3. Finally:

$$R_v(\bar{\rho}) = \ell_v(\bar{\rho}) S_v(\bar{\rho}).$$

We obtain the likelihood of interest (marginalising the prevalence and given a tree) by enforcing the constraint that the sum of all clonal prevalence is one:

$$\int_{\Delta_{|V|}} p(\boldsymbol{\rho}|\boldsymbol{\kappa}) \prod_{n=1}^{N} f(x_n|\bar{\rho}_{v_n}) d\boldsymbol{\rho} = R_r(1),$$

where $v = r$ is the root of the tree.
Pseudocode for the recursion is provided in Algorithm 1. The asymptotic runtime of this algorithm is as follows:

$$\mathcal{O}(V(NL + CSL^2))$$

Where $V$ is the number of nodes in $T_v$, $N$ is the number of mutations, $L$ is the grid size, $C$ is the number of child nodes to $v$, and $S$ is the number of samples.

##### 1.5.1 Justification

The algorithm is based on the following decomposition:

$$\begin{aligned} R_v(\bar{\rho}) &= \ell_v(\bar{\rho}) \times S_v(\bar{\rho}) \\ S_v(\bar{\rho}) &= \sum_{\bar{\rho}' \in \Phi : \bar{\rho}' \leq \bar{\rho}} (\bar{\rho} - \bar{\rho}')^{\kappa - 1} \times D_v^{|C_v|}(\bar{\rho}') \\ D_v^{|C_v|}(\bar{\rho}') &= \sum_{\{\bar{\rho}_{v_1}, \ldots, \bar{\rho}_{v_{|C_v|}} : \sum_k \bar{\rho}_{v_k} = \bar{\rho}'\}} \prod_{k=1}^{|C_v|} R_{v_k}(\bar{\rho}_{v_k}). \end{aligned}$$

In this decomposition, $\bar{\rho}'$ can be interpreted as the left hand side of the constraint imposed by the domain of integration $\mathcal{C}_T$ over the prevalences of the children:

$$\underbrace{\sum_{k=1}^{|C_v|} \bar{\rho}_{v_k}}_{\bar{\rho}'} \leq \underbrace{\bar{\rho}_v}_{\bar{\rho}}$$

#### 1.6 Inference

To simultaneously infer the posterior distribution of $T, \mathbf{b}$ we use a sequential Monte Carlo (SMC) algorithm. The SMC algorithm works by evolving a set of particles which are iteratively extended and reweighed, such that the final iteration yields an approximation of the posterior. We assume that a permutation, $\sigma$, of $[N]$ has been given. At algorithmic time $t$ of the SMC procedure we add data point $X_{\sigma(t)} = x_t$. We will grow the rooted forest from the bottom up. Because we use a fixed order of the data points, not all trees will be accessible using this procedure. Specifically, if $x = \sigma_i$ and $y = \sigma_j$ where $i < j$ then any tree where $x$ is an ancestor of $y$ cannot be reached. In the next sections we describe the basic SMC procedure, and later describe how to fix the ordering problem using a Particle Gibbs (PG) sampler.

##### 1.6.1 Target density

Let $T_t$ be the rooted tree generate at algorithmic time $t$ by connecting all existing root nodes in the forest to a dummy root. We let our target density be

$$\begin{aligned}\gamma_t(x_{1:t}) &= p(X_t, \mathbf{b}_t, T_t) \\ &= p(\mathbf{b}_t)p(T_t|\mathbf{b}_t)p(X_t|T_t)\end{aligned}$$

If $|\mathbf{b}_t| = |\mathbf{b}_{t-1}|$, we allocate a data point to an existing cluster $b$ and

$$\frac{\gamma_t(x_{1:t})}{\gamma_{t-1}(x_{1:t-1})} = |b|\frac{p(X_t|T_t)}{p(X_{t-1}|T_{t-1})}$$

otherwise we create a new cluster and

$$\frac{\gamma_t(x_{1:t})}{\gamma_{t-1}(x_{1:t-1})} = \alpha \times \frac{(|\mathbf{b}_{t-1}|+1)^{|\mathbf{b}_{t-1}|-1}}{(|\mathbf{b}_{t-1}|+2)^{|\mathbf{b}_{t-1}|}} \times \frac{p(X_t|T_t)}{p(X_{t-1}|T_{t-1})}$$

It can be checked that

$$\begin{aligned}\gamma(x_1)\prod_{t=2}^{N}\frac{\gamma_t(x_{1:t})}{\gamma_{t-1}(x_{1:t-1})} &= \frac{\alpha^{|\mathbf{b}|}\prod_{b\in\mathbf{b}}(|b|-1)!}{(|\mathbf{b}|+1)^{|\mathbf{b}|-1}}p(X|T) \\ &= p(X, \mathbf{b}, T)\end{aligned}$$

##### 1.6.2 Proposal function

To propose a new state at time $t$ we consider all possible states which can be reached by

1. Adding data point to an existing node
2. Creating a new node choosing a possibly empty subset of children from the existing root nodes.

There are several ways in which to perform these action. The simplest approach is to randomly choose to perform steps 1 or 2, and then randomly sample a node if we choose 1 or randomly select a set of children if we choose 2. While computationally inexpensive, this approach will tend to propose very poor trees and require many particles.

Alternatively we could enumerate all possible choices and compute the probability of the trees which result. We can then sample proportional to these probabilities. This approach becomes computationally expensive as we need to enumerate all possible subsets of children if we start a new node.

In practice we use a hybrid approach. We randomly choose whether to join an existing node or start a new node. When adding a mutation to an existing node we compute the probability of the resulting tree with the mutation attached at each node. When adding a mutation to a new node we randomly sample the set of children. This approach provides a compromise between the quality of trees proposed and the computational complexity.

##### 1.6.3 Particle Gibbs sampler

We treat the ordering of data points added at algorithmic time $t$ as fixed before running the SMC procedure. This limits the utility of the SMC algorithm, as only a subset of trees can be reached. For example, the first element of $\sigma$ can never be a root node unless the forest has only a single node. To address this issue we use a particle Gibbs (PG) sampler to embed our SMC procedure in a more general Markov chain Monte Carlo sampler. With a slight abuse of notation we will use $p(X, T)$ to denote $p(X, \boldsymbol{b}, T)$ where it understood each node in the tree is associated with a cluster of mutations. We define $\Sigma(T)$ to be the set of permutations which could generate $T$ using the SMC inference scheme. Instead of constructing an SMC algorithm to target the posterior $p(T|X)$ directly, we alternate between using conditional SMC to target $p(T|X, \sigma)$ and sampling from $p(\sigma|T)$.

$$
\begin{aligned}
p(T|X,\sigma) &\propto p(T,X,\sigma) \\
&= p(\sigma|T)p(X|T)p(T) \\
p(\sigma|T) &= \frac{\sigma \in \Sigma(T)}{\sum_\sigma \sigma \in \Sigma(T)} \\
&= \frac{\sigma \in \Sigma(T)}{Z(\sigma)}
\end{aligned}
$$

A $\sigma$ will be in $\Sigma(T)$ if it could be generated by the following recursive algorithm.

1. For a leaf node, return a randomly shuffled list of its assigned mutations.
2. For an internal node, recursively collect the sigma lists returned from processing its child nodes. Interleave these child-node lists using a bridge shuffle to create a flat sigma list; then collect the datapoints assigned to the current node, shuffle their order, and append them to the end of the current sigma list.
3. At the root node, proceed as above in the internal node case, with the added step of also collecting the outlier datapoints (should any exist), shuffling their order and interleaving the outliers with the sigma list.

###### 1.6.4 Particle Gibbs sub-tree sampling

The above procedure can be easily modified to update sub-trees instead of the whole tree. This can alleviate the well known degeneracy problem for PG samplers. Degeneracy refers to the problem that for long sequences all particles will eventually trace their genealogy back to the conditional path in the swarm. This limits the ability of the sampler to change states that appear early in the SMC iterations. In our case this means that deeper nodes in the tree will tend to change less frequently. By re-sampling sub-trees we can shorten the SMC paths and more efficiently update deeper nodes in the tree.

To select a sub-tree for updating, we randomly sample a mutation and find the node the mutation is assigned to. We then take the parent of that node as the root of the sub-tree which we will update. Next we use the PG sampler described above to update sub-tree. At the final iteration we reinsert the sub-tree into the original tree and compute the full tree likelihood to use in the computation of the importance weights. This ensures the final target density in the conditional SMC is proportional to the target posterior distribution.

One non-obvious aspect of the above algorithm is the mechanism for choosing the node to use as the sub-tree root. We choose a mutation at random so that the update does not depend on the current state of the Monte Carlo sampler. We choose to use the parent of the associated node as the sub-tree root in order to allow the sampler to update the entire forest. If we did not do this, we could never change the attachments to the dummy root.

##### 1.7 Outliers

A major constraint of the model described thus far is that mutations are assumed to be propagated to all nodes that descend from the node of origin. This constraint can easily be violated if copy number alterations remove the mutation in a descendant. Furthermore, we may have mutations which may be noisy either due to erroneous copy number or incorrect allele counts due to alignment errors. To address these issues the above model can be extended with an outlier state. Each mutation is assigned a prior probability of being an outlier, $\nu_n$. We define a binary variable $o_n$ which indicates whether mutation $n$ is an outlier. If a mutation is not an outlier it joins a node in the tree and contributes to the likelihood as above. If the mutation is an outlier it has the standard PyClone likelihood where we assume it has a cellular prevalence drawn from a uniform distribution. We marginalise the cellular prevalence in this case using numerical integration. The updated joint likelihood becomes

$$
\begin{aligned}
p(X, \mathbf{b}, T, \mathbf{o}) &= \prod_{n=1}^{N} [\nu_n \int f(x_n|\rho) d\rho]^{\mathbb{I}(o_n=1)} \\
&\times \prod_{n=1}^{N} p(\mathbf{b}|\alpha) p(T|\mathbf{b}) \int p(\boldsymbol{\rho}|\boldsymbol{\kappa}) \\
&\times \prod_{n=1}^{N} [(1-\nu_n) f(x_n|\bar{\rho}_{v_n})]^{\mathbb{I}(o_n=0)} d\boldsymbol{\rho}
\end{aligned}
$$

The modifications to the inference procedure are straightforward. When proposing a new state we now include the possibility that a mutation is an outlier. When re-sampling $\sigma$ we randomly permute the set of outliers and interleave them with the permuted values from the tree using a bridge shuffle. The algorithm for counting the number of permutations can be easily modified to account for this. Finally, when performing sub-tree updates we include all outliers in the set of mutations to be considered for the sub-tree update.

##### 1.7.1 Outlier prior probability assignment from clustered data

PhyClone can assign prior outlier probabilities for clusters given a clustered input file which includes: SNV genomic position (chromosome and position), and SNV or cluster level cellular prevalence information. Given the appropriate input, PhyClone will, in a manner not dissimilar from that employed by CONIPHER [1], assign clusters with either a high or low outlier prior probability (both levels are user-defined, default to 0.01 for low and 0.5 for high outlier prior probability).

The first stage in the data informed approach is to first identify the cluster most likely to be truncal in the phylogeny, this is achieved by selecting the cluster with the highest cellular prevalence across samples. Should there be multiple candidate clusters from this primary filtering, a second stage is employed where the mean of all cellular prevalence across all samples are computed for each candidate cluster; the truncal cluster will then be the cluster with the maximum mean cellular prevalence from the truncal candidates.

Following truncal cluster selection, a background distribution of mutations per unique chromosome is established from the truncal cluster. For each other cluster, given its set of mutations $m$, will have its expected number of unique chromosomes computed by making 10,000 random draws of size $|m|$ from the truncal background distribution. From these 10,000 random trials, the number of trials with fewer unique chromosomes than what is observed ($N_u$) will be used to compute the resulting p-value ($\frac{N_u}{10000}$). Clusters with a computed $p$-value $< 0.01$ will be defined as more likely to be lost due to a closer than expected genomic locality.

#### 2 Supplementary Figures

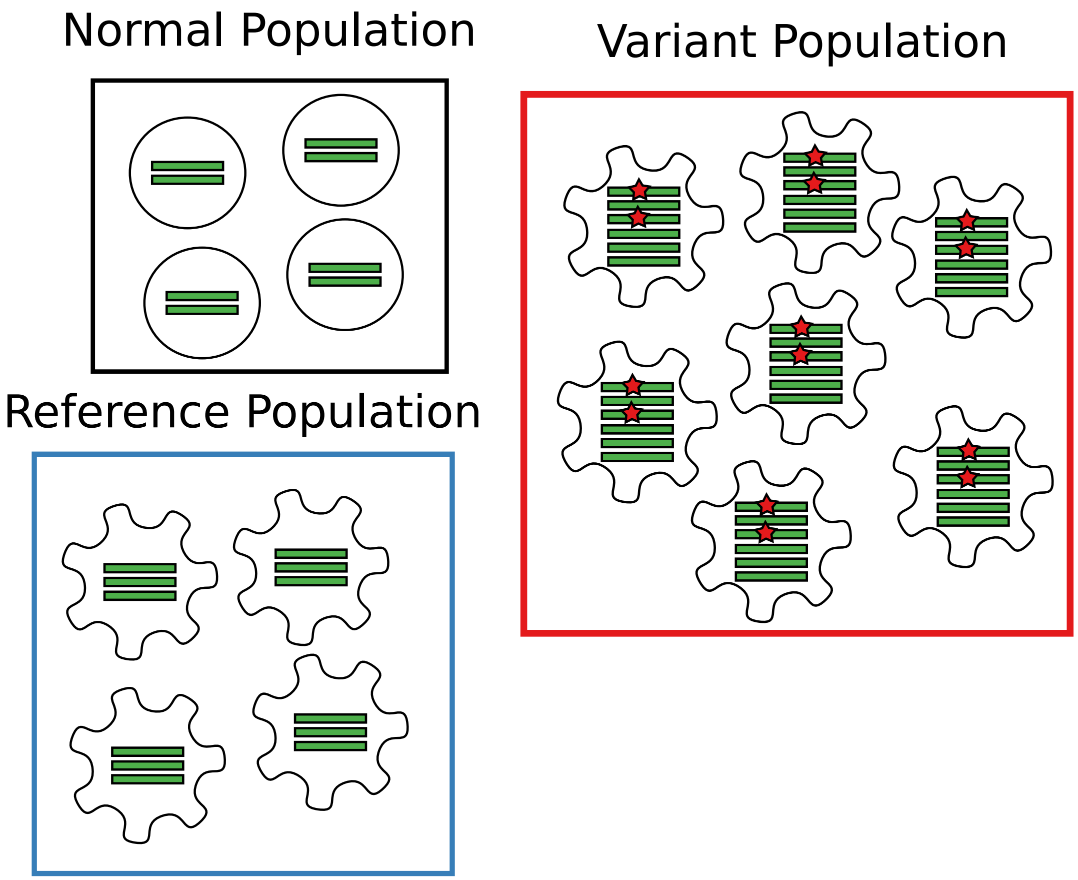


Figure 1: **Population structure assumed by the PyClone emission density**. Circular shaped cells are healthy cells, while irregular shaped cells are cancerous. The green bars are indicate chromatids and the red stars indicate a mutation. These are all assumed to overlap the loci which contains the mutation.

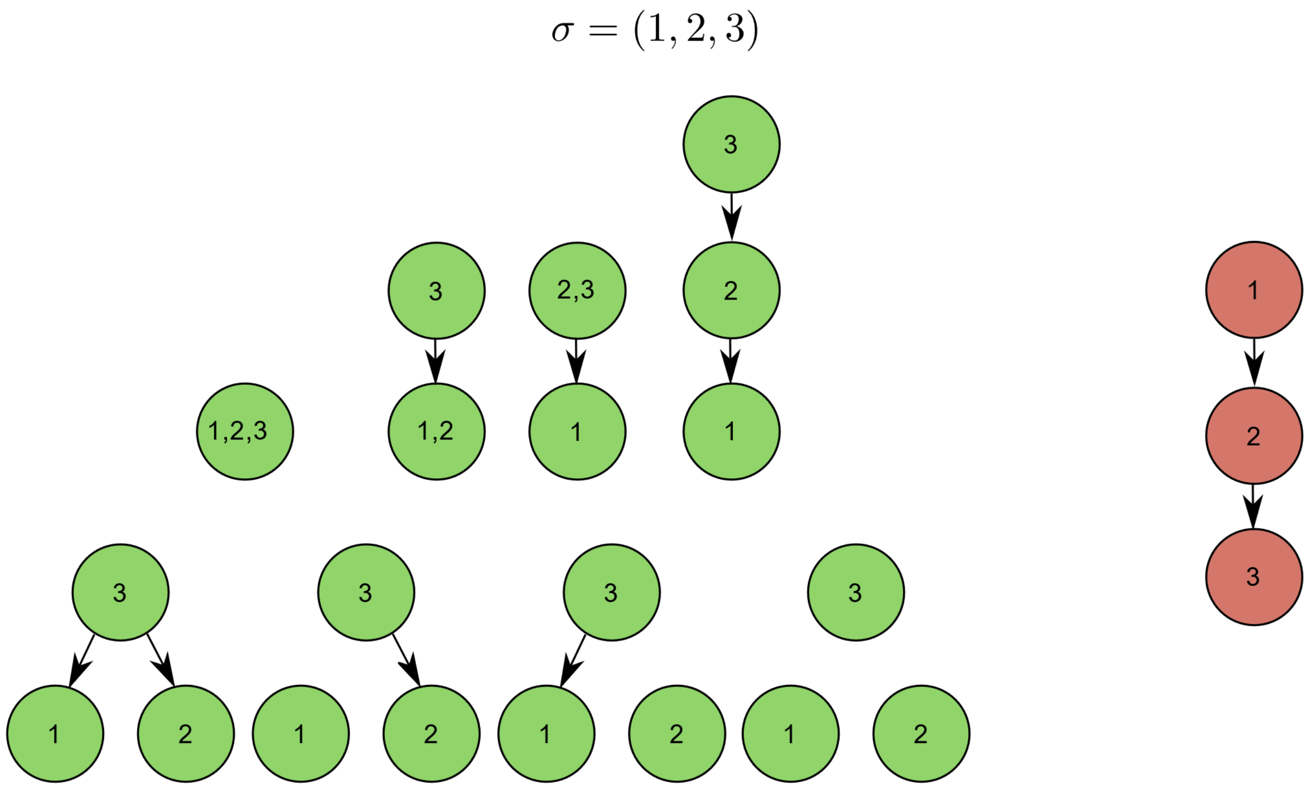


Figure 2: **SMC, Not all trees are reachable**. Graphical representation of how the data point ordering given to an SMC method can impact its end result when building up from the root. In this figure the tree on the right (in red), is unreachable by pure SMC due to the given data ordering; while all of the topologies on the left (in green), represent the topologies reachable by a pure SMC approach with the given data ordering.

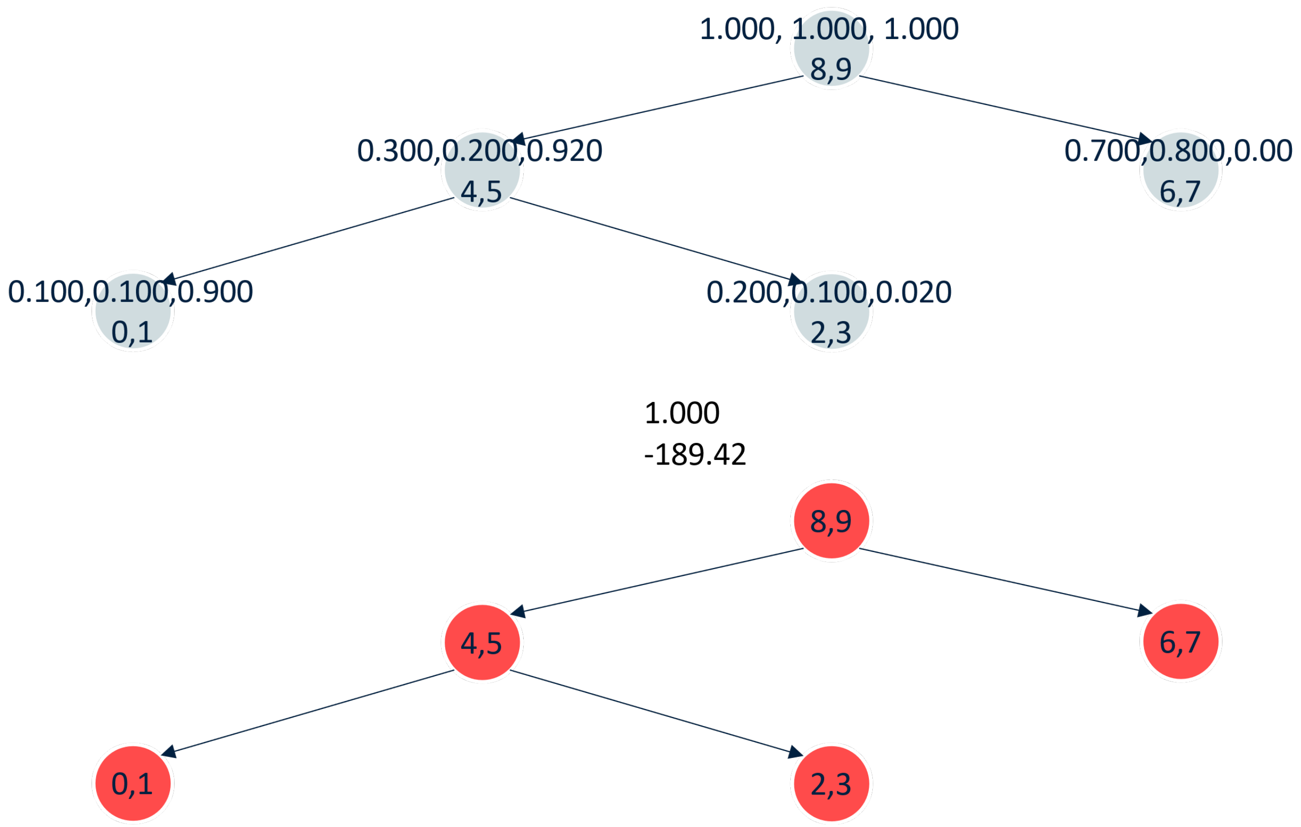


Figure 3: **SMC given a good data ordering**. Graphical representation of how the data point ordering given to an SMC method can impact its end result when building up from the root. Data ordering given as $\sigma = (0, 1, \ldots, 8, 9)$. The upper tree (in blue), represents the ground truth topology, while the lower tree (in red) represents the result from a pure SMC approach to building the topology with the given $\sigma$.

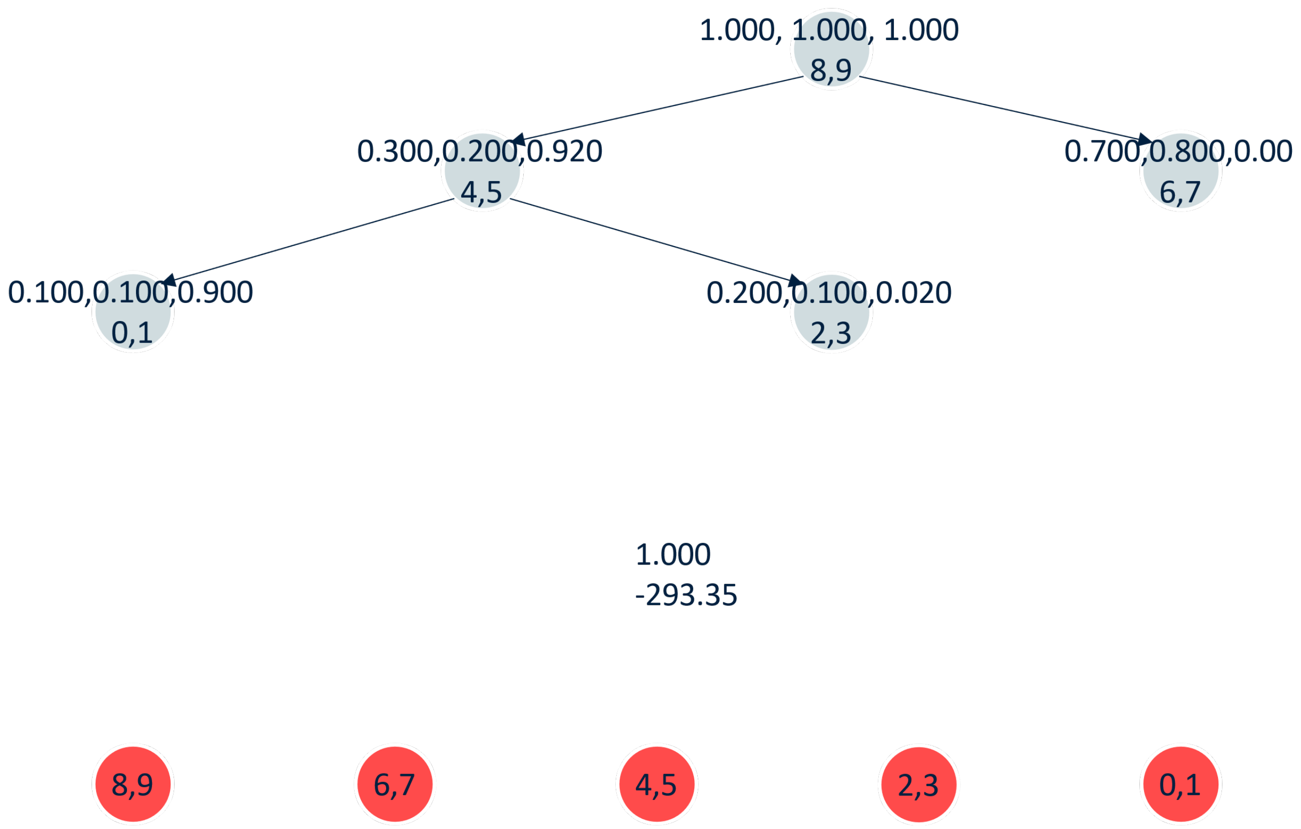


Figure 4: **SMC given a bad data ordering**. Graphical representation of how the data point ordering given to an SMC method can impact its end result when building up from the root. Data ordering given as $\sigma = (9, 8, \ldots, 1, 0)$. The upper tree (in blue), represents the ground truth topology, while the lower tree (in red) represents the result from a pure SMC approach to building the topology with the given $\sigma$.

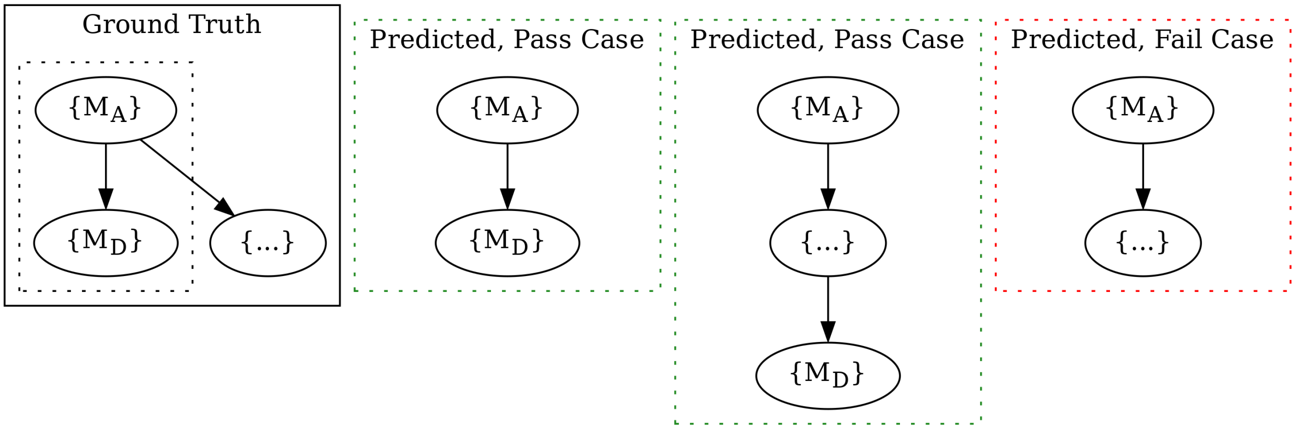


Figure 5: **Ancestor-Descendant reconstruction validity**. Graphical representation of how the ancestor-descendant relation metric defines a valid reconstruction. Curly brackets "{}" in nodes represent the set of SNVs assigned to that node. Ground truth box represents the true ancestor-descendant relationship between a single pair of SNVs $\{M_A, M_D\}$, where $M_A$represents an SNV that has an ancestor relationship to $M_D$.

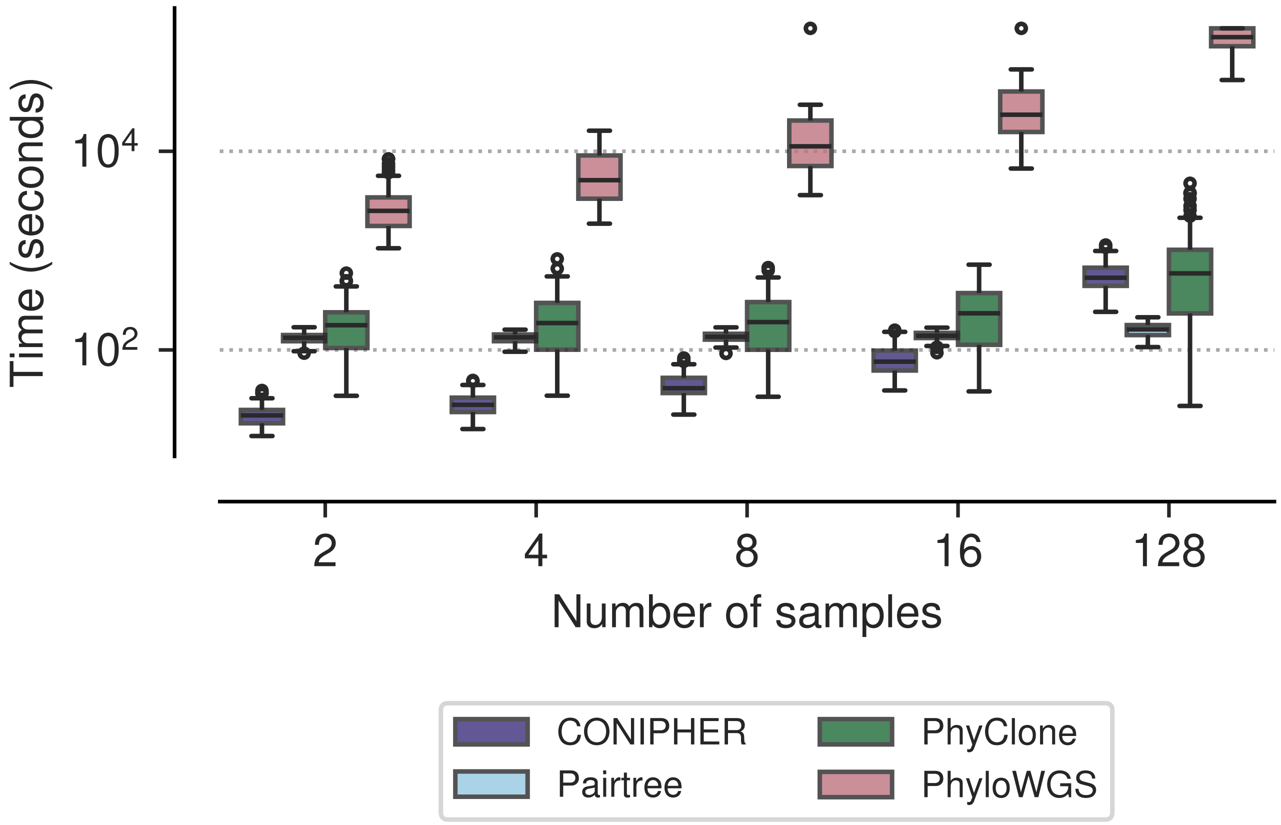


Figure 6: **Running time on small TSSB dataset by number of samples**. Running time in seconds for each method to analyse trials from the small TSSB dataset.

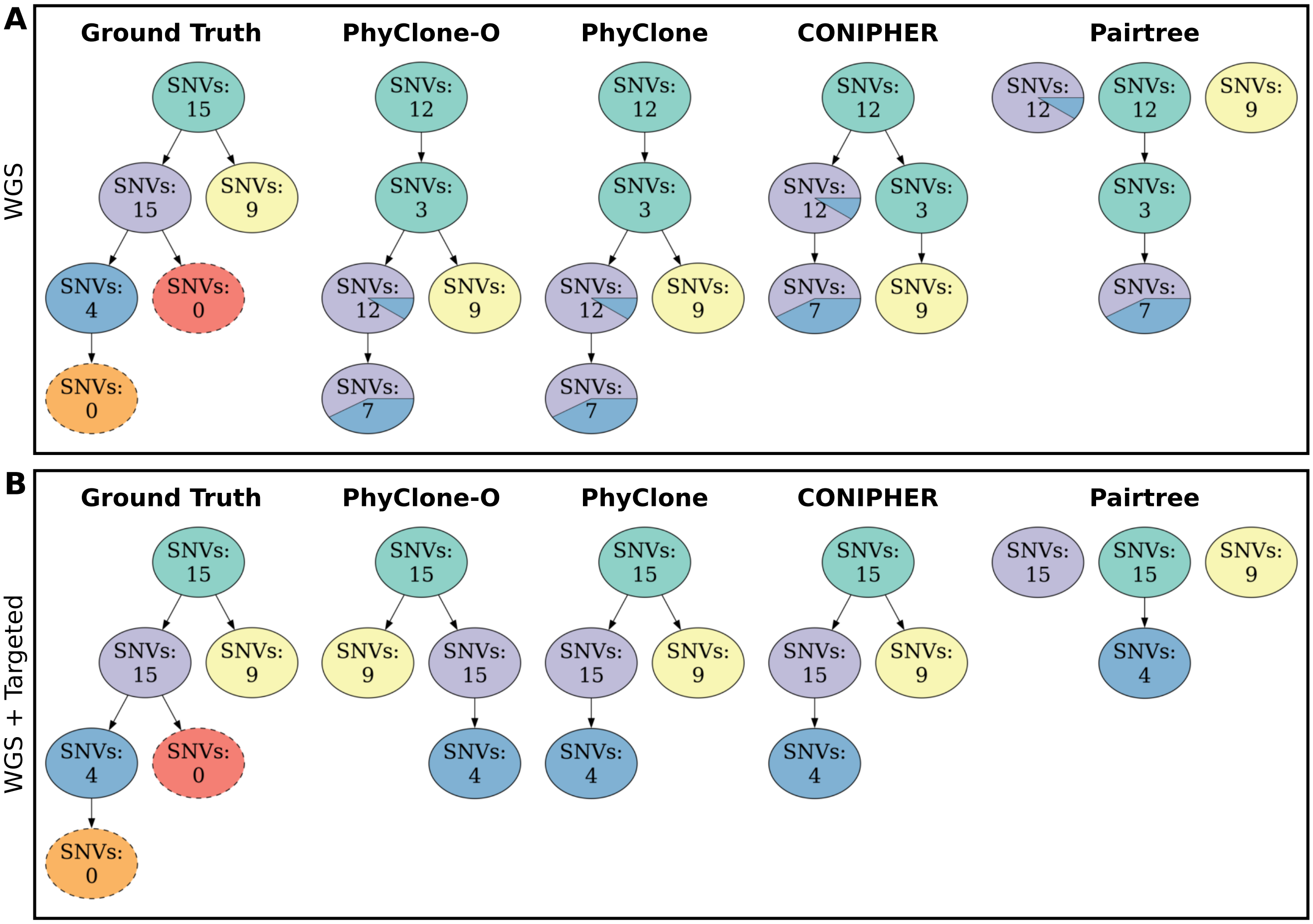


Figure 7: **HGSOC patient 2 clonal phylogenetic trees**. **A)** Predicted trees built using purely Whole Genome Sequencing (WGS) data. **B)** Predicted trees built using a combination of WGS and targeted deep sequencing data [2]. **i**) Ground truth phylogenetic tree inferred from single cell and targeted deep sequencing data [2], nodes with a dashed border denote clones that are defined only by the absence of SNVs from the parent. Predicted trees built using: **ii**) PhyClone with outlier modelling; **iii**) PhyClone without outlier modelling; **iv**) CONIPHER; and **v**) Pairtree. Colours of nodes in method inferred trees correspond to the single nucleotide variant clonal (SNV) assignment from ground truth.

**Algorithm 1** PhyClone Marginalisation

**Input:** $T = (V, E)$, $\bar{\boldsymbol{\rho}} = (\bar{\rho}_1, \ldots, \bar{\rho}_{|V|})$, $X$ = observed copy number and read count data
**Output:** Node parameter marginalised likelihood of $T$

**procedure** TREE-LIKELIHOOD-MARGINALIZATION($T$, $\bar{\boldsymbol{\rho}}$, $X$)
    **for** $v \in V$, post-order from $T$ **do**
        $T_v \Leftarrow (V_v, E_v)$
        $R_v \Leftarrow$ COMPUTE $R(T_v, \bar{\boldsymbol{\rho}}_v)$
    **end for**
**end procedure**

**function** COMPUTE $R(T_v, \bar{\boldsymbol{\rho}}_v)$
    $\ell_v \Leftarrow$ COMPUTE $\ell_v(\bar{\boldsymbol{\rho}}_v)$
    $C_v \Leftarrow$ child nodes of $v$
    **if** $|C_v| = 0$ **then**
        **return** $\ell_v$
    **else**
        $S_v \Leftarrow$ COMPUTE $S(C_v)$
        **return** $\ell_v \times S_v$
    **end if**
**end function**

**function** COMPUTE $S(C_v)$
    $D \Leftarrow$ COMPUTE $D(C_v)$
    $S[1] \Leftarrow D[1]$
    **for** $i \in (2, \ldots, |D|)$ **do**
        $S[i] \Leftarrow D[i-1] + D[i]$
    **end for**
    **return** $S$
**end function**

**function** COMPUTE $D(C_v)$
    $D \Leftarrow R_{C_v[1]}$
    **for** $k \in (2, \ldots, |C_v|)$ **do**
        $D \Leftarrow D * R_{C_v[k]}$
    **end for**
    **return** $D$
**end function**

**function** COMPUTE $\ell_v(\bar{\rho})$
    $\ell_v \Leftarrow 1$
    **for** $n \in v$ **do**
        $\ell_v \Leftarrow \ell_v \times$ PYCLONE$(x_n | \bar{\rho}_{v_n})$
    **end for**
    **return** $\ell_v$
**end function**

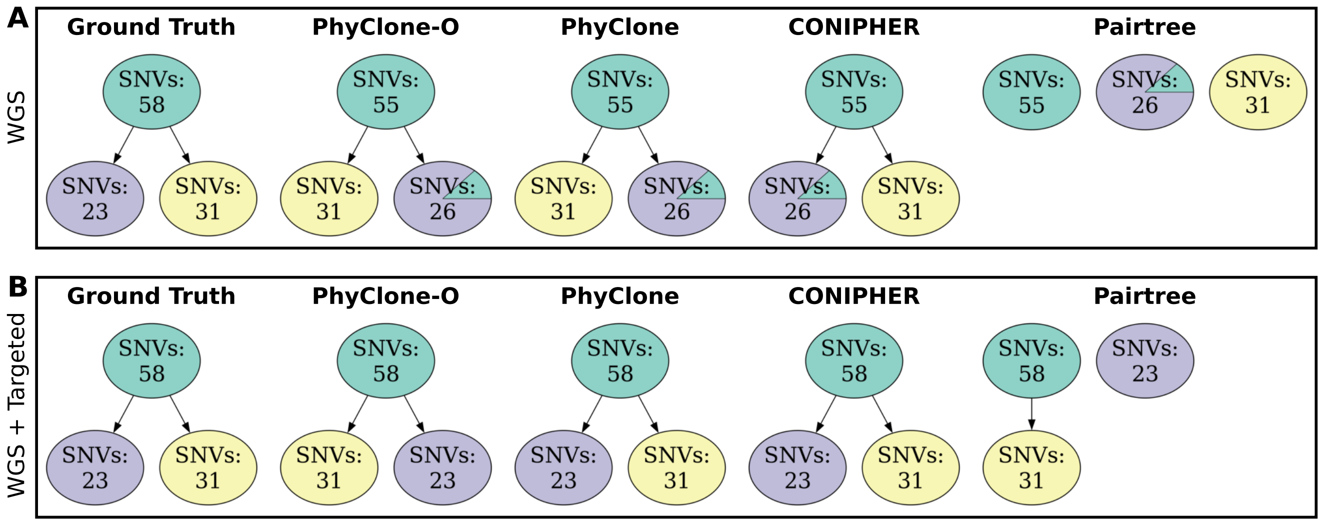


Figure 8: **HGSOC patient 9 clonal phylogenetic trees**. **A)** Predicted trees built using purely Whole Genome Sequencing (WGS) data. **B)** Predicted trees built using a combination of WGS and targeted deep sequencing data [2]. **i**) Ground truth phylogenetic tree inferred from single cell and targeted deep sequencing data [2] nodes with a dashed border denote clones that are defined only by the absence of SNVs from the parent. Predicted trees built using: **ii**) PhyClone with outlier modelling; **iii**) PhyClone without outlier modelling; **iv**) CONIPHER; and **v**) Pairtree. Colours of nodes in method inferred trees correspond to the single nucleotide variant clonal (SNV) assignment from ground truth.

#### 3 Supplementary Tables

Table 1: **High grade serous ovarian cancer data performance metrics**. Reported metrics were computed from predicted phylogenetic trees from two datasets, purely Whole Genome Sequencing (WGS) data, and a combination of WGS and targeted deep sequencing (W+T) data.

| Program | Patient | V-Measure | | AD F-Score | |
|---|---|---|---|---|---|
| | | WGS | W+T | WGS | W+T |
| PhyClone | 9 | 0.91 | 1.00 | 0.95 | 1.00 |
| PhyClone-O | 9 | 0.91 | 1.00 | 0.95 | 1.00 |
| CONIPHER | 9 | 0.91 | 1.00 | 0.95 | 1.00 |
| Pairtree | 9 | 0.91 | 1.00 | 0.00 | 0.73 |
| PhyClone | 2 | 0.78 | 1.00 | 0.89 | 1.00 |
| PhyClone-O | 2 | 0.78 | 1.00 | 0.89 | 1.00 |
| CONIPHER | 2 | 0.78 | 1.00 | 0.82 | 1.00 |
| Pairtree | 2 | 0.78 | 1.00 | 0.34 | 0.22 |
| PhyClone | 3 | 0.73 | 0.86 | 0.69 | 0.82 |
| PhyClone-O | 3 | 0.85 | 1.00 | 0.82 | 0.93 |
| CONIPHER | 3 | 0.73 | 0.86 | 0.70 | 0.80 |
| Pairtree | 3 | 0.73 | 0.86 | 0.42 | 0.53 |

**Supplementary Tables S1–S13**: Benchmarking Performance Result Metrics

- **S1**: TSSB small
- **S2**: TSSB large
- **S3**: FS-CRP loss
- **S4**: Pairtree Low-Complexity (LC)
- **S5**: Pairtree High-Complexity (HC)
- **S6**: CONIPHER No-Noise
- **S7**: CONIPHER with noise
- **S8**: HGSOC WGS, patient 2
- **S9**: HGSOC WGS + Targeted, patient 2
- **S10**: HGSOC WGS, patient 3
- **S11**: HGSOC WGS + Targeted, patient 3
- **S12**: HGSOC WGS, patient 9
- **S13**: HGSOC WGS + Targeted, patient 9

**Supplementary Tables S14–S26**: Benchmarking Friedman-Nemenyi Analysis Results

- **S14**: CONIPHER No-Noise Friedman results
- **S15**: CONIPHER No-Noise Nemenyi results

- **S16**: CONIPHER with noise Friedman results
- **S17**: CONIPHER with noise Nemenyi results
- **S18**: FS-CRP loss Nemenyi results
- **S19**: Pairtree High-Complexity (HC) Friedman results
- **S20**: Pairtree High-Complexity (HC) Nemenyi results
- **S21**: Pairtree Low-Complexity (LC) Friedman results
- **S22**: Pairtree Low-Complexity (LC) Nemenyi results
- **S23**: TSSB large Friedman results
- **S24**: TSSB large Nemenyi results
- **S25**: TSSB small Friedman results
- **S26**: TSSB small Nemenyi results

### References

[1] Kristiana Grigoriadis, Ariana Huebner, Abigail Bunkum, Emma Colliver, Alexander M. Frankell, Mark S. Hill, Nicolai J. Birkbak, Charles Swanton, Simone Zaccaria, and Nicholas McGranahan. Conipher: A computational framework for scalable phylogenetic reconstruction with error correction. *Protocol Exchange*, feb 2023.

[2] Andrew McPherson, Andrew Roth, Emma Laks, Tehmina Masud, Ali Bashashati, Allen W Zhang, Gavin Ha, Justina Biele, Damian Yap, Adrian Wan, Leah M Prentice, Jaswinder Khattra, Maia A Smith, Cydney B Nielsen, Sarah C Mullaly, Steve Kalloger, Anthony Karnezis, Karey Shumansky, Celia Siu, Jamie Rosner, Hector Li Chan, Julie Ho, Nataliya Melnyk, Janine Senz, Winnie Yang, Richard Moore, Andrew J Mungall, Marco A Marra, Alexandre Bouchard-Côté, C Blake Gilks, David G Huntsman, Jessica N McAlpine, Samuel Aparicio, and Sohrab P Shah. Divergent modes of clonal spread and intraperitoneal mixing in high-grade serous ovarian cancer. *Nature Genetics*, 48(7):758–767, may 2016.

[3] Andrew Roth, Jaswinder Khattra, Damian Yap, Adrian Wan, Emma Laks, Justina Biele, Gavin Ha, Samuel Aparicio, Alexandre Bouchard-Côté, and Sohrab P Shah. PyClone: statistical inference of clonal population structure in cancer. *Nature Methods*, 11(4):396–398, mar 2014.
